# Temporal Profiling of Upper-Layer Neurons Reveals Changes in the Molecular Landscape Upon Maternal Immune Activation

**DOI:** 10.64898/2026.04.14.718455

**Authors:** Tabitha Rücker, Birgit Schwanke, Kerstin Reumann, Patrick Blümke, Jacqueline Nakel, Daniela Indenbirken, Thomas Günther, Thomas Mair, Bente Siebels, Paula Nissen, Kilian Müller, Melanie Richter, Sabine Hoffmeister-Ullerich, Stephanie Stanelle-Bertram, Gülsah Gabriel, Adam Grundhoff, Hartmut Schlüter, Froylan Calderon de Anda

**Author notes:** Present address: Epigenomic, Proliferation and Cell Identity Unit, Institut Pasteur, 25, rue du Docteur Roux, 75015 Paris, France (Tabitha Rücker). To whom correspondence should be addressed: Tabitha Rücker, Froylan Calderón de Anda.

## Abstract

Neuronal differentiation is a dynamic, multi-layered process that transforms progenitor cells into functionally integrated neurons, yet the coordinated molecular programs underlying this transition remain incompletely understood. Here, we define the developmental trajectory of murine upper-layer cortical neurons using an integrated multi-omics approach combining transcriptomics and proteomics across key stages of neurogenesis.

We find that neuronal identity emerges gradually through coordinated transitions from RNA processing and splicing programs toward synaptic and metabolic maturation. Integration of matched transcriptomic and proteomic datasets reveals a compact set of concordant molecular modules that define this trajectory, highlighting post-transcriptional regulation as a central driver of neuronal maturation.

We further show that maternal immune activation (MIA), a model of prenatal inflammation, deranges this developmental program. MIA induces sustained upregulation of Wnt signalling pathways alongside a downregulation of synaptic regulators, without detectable global alterations in DNA methylation. These molecular changes are accompanied by defects in neuronal positioning during cortical development, linking altered molecular trajectories to functional outcomes.

Together, our findings establish a temporal molecular framework of neuronal differentiation and demonstrate that prenatal environmental perturbations reshape cortical development primarily through post-transcriptional and signalling-based mechanisms.

## Introduction

Neuronal development is a complex, finely tuned process that underlies the formation of a functional brain. Once generated, neurons are traditionally believed to exit the cell cycle permanently and differentiate irreversibly, a process that is maintained throughout their lifespan^1–3^. The precise molecular mechanisms governing their developmental transition from a progenitor cell to the initiation of neuronal connectivity are not fully understood^4^. Unravelling these mechanisms is paramount, as disturbances in neuronal development can lead to significant neurological and cognitive impairments later in life.

The cerebral cortex, particularly its upper layers, plays a crucial role in higher-order brain functions such as sensory perception, cognition, and motor control^5^. Moreover, upper-layer cortical excitatory neurons are most dysregulated in the cortex of idiopathic ASD patients^6^. Understanding the development of upper-layer cortical neurons (ULNs) is therefore critical for elucidating how the brain processes complex information.

In this study, we analyse the lineage trajectory of ULNs using external information from real-time follow-up of developing neurons, without relying on marker genes defined *a priori*, enabling us to examine their trajectory in transcriptomic, proteomic, and morphological space. Our comprehensive characterisation spans from their generation at embryonic day 14 (E14) to postnatal day 7 (P7), when their synaptic connections extend to the other hemisphere. To ensure broad coverage of timely, fine-tuned processes, we isolated specific cell populations daily, including neuronal progenitors, early-differentiating neurons, and a mixed cell population. These cell populations were labelled *in utero* with fluorescent dyes driven by the Glast1, NeuroD1, and a non-specific CAG promoter with high constitutive expression. We then employ a multimodal analytical approach that integrates transcriptomic and proteomic analyses^7^. This strategy enabled the capture of dynamic molecular signatures underlying the developmental trajectory of a defined population of neurons (ULNs). Our results highlight that the neuronal identity of early-developing cortical ULNs is acquired gradually, as we previously suggested^8^. Our results indicate a cell fate transition based on the downregulation of progenitor-associated genes and a highly specific upregulation of the unique neuronal transcriptomic landscape at the birth stage. In addition, this approach reveals differences between the transcriptome landscape of cells isolated with a non-specific versus a neuronal-specific promoter; thus, underscoring the relevance of our method for the longitudinal analysis of defined developing neuronal populations^7^ compared to previously reported pulse-labelling approaches^9,10^.

Extrinsic factors, particularly those encountered prenatally, can profoundly influence neuronal development. One such factor is maternal immune activation (MIA), which models the effects of maternal infection and inflammation and is implicated in altering brain development and increasing the risk of neurodevelopmental disorders in offspring^11–15^. Our findings reveal that MIA exposure leads to an upregulation of Wnt signalling pathways and a concurrent downregulation of key synaptic transcripts without altering the methylome landscape of developing neurons upon MIA exposure. These changes in key molecular pathways suggest that MIA can hinder normal neuronal development and maturation, potentially impacting brain connectivity and function. As these changes are primarily post-transcriptional, this highlights potential targets for therapeutic strategies to mitigate the detrimental effects of prenatal immune challenges on neurodevelopment.

## Results

We aimed to isolate and characterise defined ULNs during critical periods of development. To this end, the commitment to an upper-layer neurogenic fate by cortical progenitor cells was investigated using a multimodal approach that combined flow cytometric isolation of defined cell populations across various developmental time points, as previously described^7^. Radial glial cells (RGCs) were labelled via *in utero* electroporation (IUE) at E12 and isolated at E14, while early postmitotic neurons destined for upper layers were labelled at E14 and harvested from E16 through P7 (**Fig. 1a; Supp. Fig. 1a**). Sorted populations from matched biological samples were subject to bulk proteomics, RNA sequencing, and Enzymatic Methylation (EM)-seq (**Fig. 1a, b**). E19 samples were separated into yet unborn embryos (E19_E) and postnatal pups (E19_P) but were combined into a single group for proteomic analysis, due to limited availability (**Supp. Fig. 1b**).

**Figure 1.**
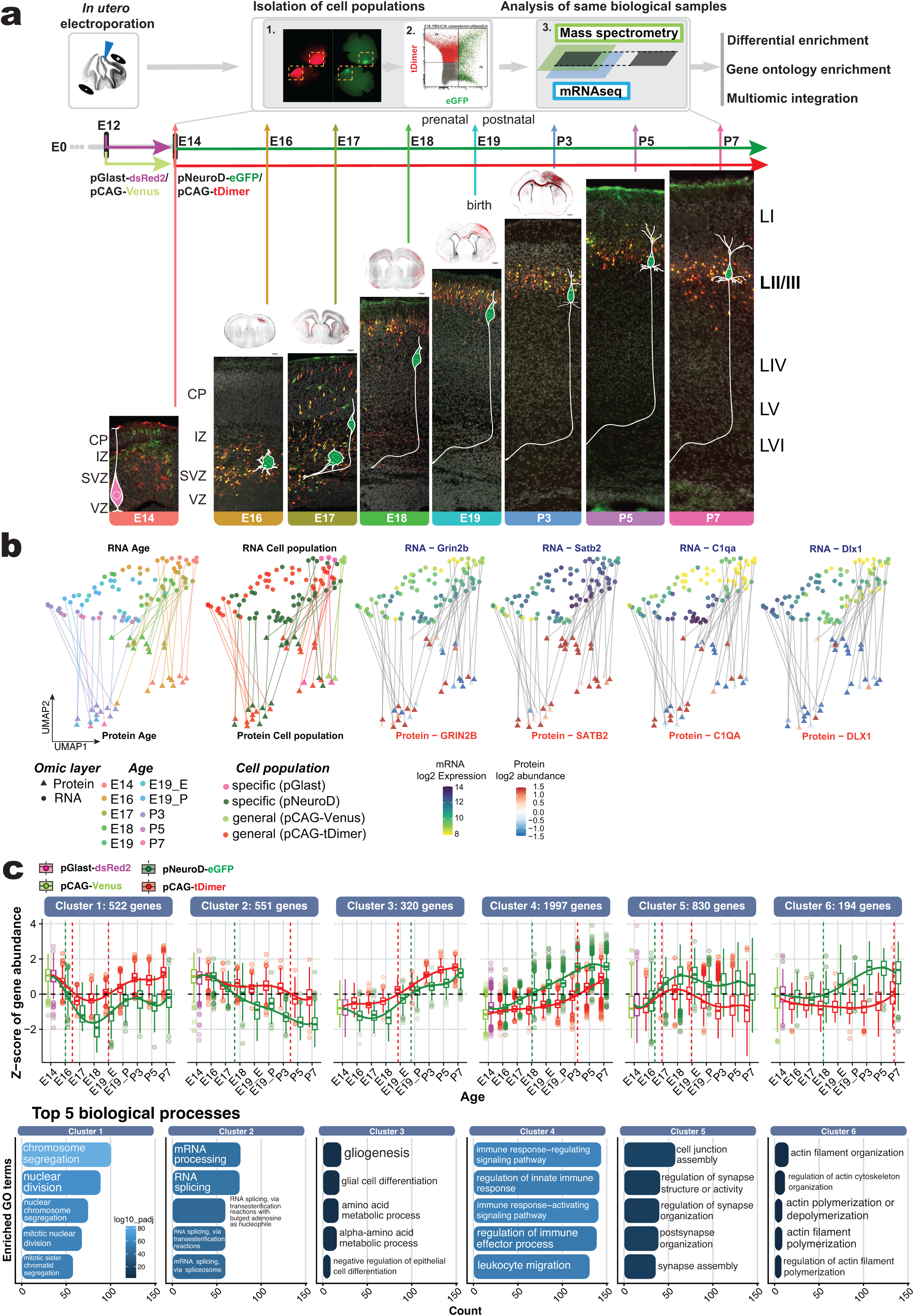
Experimental overview to target and study molecular changes of the developing upper-layer neurons. **a)** *Experimental strategy for the temporal dissection of cortical layer II/III neurogenesis. In utero* electroporation (IUE) at E14 labels cortical progenitors, which were isolated by FACS and processed for RNA-seq/EM-seq or proteomic mass spectrometry, followed by integrative bioinformatic analysis. **Bottom:** Representative coronal sections showing fluorophore expression from E14 to P7. For E14-P3, 4× overviews (Hoechst, grey; tDimer, red) are shown above higher-magnification images. Scale bar, 100 µm. **b)** *Overview of the used samples and their features for the following analyses.* Separate UMAP embeddings of RNA-seq and protein abundance data depict biological samples across embryonic age (E14-P7) and cell population identity (circles, RNA; triangles, protein). Grey lines connect matched samples across modalities. Expression of representative markers (Satb2, C1qa, Grin2b, Dlx1) illustrates age-dependent and population-specific regulation. **c)** *Six DEG clusters highlight differences between the pCAG+ and pGlast+/pNeuroD+ populations across development.* Compared with pCAG+ cells, pNeuroD+ cells show reduced expression of cell cycle-, RNA processing-, and gliogenesis-associated genes and increased expression of genes linked to innate immune signalling, synapse organisation, and actin cytoskeleton dynamics. Cluster extrema occur predominantly at E18, marking the onset of transcriptional divergence and neuronal commitment. *pCAG+ population, red; pGlast+/pNeuroD+ population, green*.

Our results showed that at the peak of neurogenesis (at E14 in mice) labelled pGlast1-dsRed2+ RGCs were observed in the ventricular zone (VZ) as apical progenitor cells with basal projections reaching the pial surface (**Fig. 1a**). Another part of the labelled pGlast1-dsRed2+ population accumulated at the cortical plate (CP) border without clear polarity, while some cell bodies were already in contact with the pial surface. By E16, the progenitor cells were gathering in the subventricular zone (SVZ), and more mature neuronal progenitors adopted a bipolar morphology and initiated radial migration. From this point onwards, the nascent CP thickened and postmitotic neurons were subsequently labelled with pNeuroD1-eGFP. At E17, most developing neurons attained the motile stage and crossed the subplate boundary into the CP. By E18, most radially migrating neurons reached their final position in the upper layers, while a minority had not yet completed translocation. Concurrently, the volume of the SVZ decreased markedly, the CP continued to expand, and the apically located axon bundles thickened. At birth (E19), the cortex exhibited an ordered, laminar-structured CP with neurons properly oriented within their laminar niche. Immunofluorescence staining against SATB2 confirmed the upper-layer neuronal character of the pNeuroD1-eGFP+ cells (**Supp. Fig. 1c**). From P3 onwards, these neurons displayed a pyramidal shape with elongated dendrites and an axon projecting across the *corpus callosum* to the other hemisphere (**Fig. 1a**^8^). Visually, apart from the labelled progenitors at E14, the cell populations developed similarly regardless of reporter expression (eGFP+ vs tDimer+). Overall, cortical development in our model recapitulated the established sequence of events in cortical development, showing morphological transitions from neuronal progenitors to upper-layer pyramidal cells, including polarity changes, radial migration, as well as axon and dendrite formation.

Building on these observations, we employed RNA-seq to characterise the underlying transcriptional programs. Consistent with the experimental design, transcripts from the fluorescence reporters driven by a developmentally-active promoter were readily detectable in the RNA-seq dataset, exhibiting approximately ten-fold higher expression than the mean of the control/pCAG+ population within each experiment (**Supp. Fig. 1d**). In addition, RNA-seq and RT-PCR measurements showed strong concordance, thereby validating the RNA-seq results (**Supp. Fig. 1e**).

Our RNA-seq samples clustered distinctly by developmental stage and by the reporter construct that was used to distinguish cell populations, thus reflecting the gradual progression from the progenitor to the mature state (**Supp. Fig. 1f**). Because the protein samples showed weaker clustering than the mRNA samples, we subsequently integrated the proteomic and transcriptomic data to extract the most information from the protein layer, despite its greater variability.

Our protein samples matched the biological mRNA samples used for the analysis. Uniform Manifold Approximation and Projection (UMAP) embeddings of the PCA outputs highlight those 17 matched biological pGlast/pNeuroD+ samples, which are connected by lines to indicate correspondence (**Fig. 1b**, 35 matched biological samples if including the pCAG cell population). Representative genes associated with specific cell populations and age-related patterns reveal a gradual continuum of developmental and population-dependent changes across both omic layers – sometimes aligned, sometimes diverging. For example, ***Grin2b***, a subunit of the NMDA receptor with crucial roles in excitatory synaptic function and dendritic development, has been linked to promoting neuronal differentiation and early synaptogenesis in developing neurons^16^. In line with this, both its transcripts as well as GRIN2B protein levels increased toward the postnatal stage, particularly within the pNeuroD+ population. The upper-layer neuronal marker ***Satb2***, which marks callosal projection neurons and contributes to cortical circuit formation and synaptic connectivity^17^, peaked at E18 in the pNeuroD+ population with high protein abundance. In contrast, ***C1qa***, a classical complement component implicated in microglia-mediated synaptic pruning and neural circuit refinement during development^18^, showed a steady increase in expression across developmental time; however, its protein levels do not correspond to its transcript levels (**Fig. 1b**).

To identify coordinated temporal expression changes between the pCAG vs. non-pCAG sorted cell populations, we derived time-course patterns focusing on these cell populations. This time-resolved clustering revealed 5,489 Differentially Expressed Genes (DEGs) between the two cell populations (pCAG+ vs. pGlast/pNeuroD1 cell populations) from E14 to P7 at a padj. level of <0.01, which were split into 6 clusters (**Fig. 1c**). The biological functions associated with these differential dynamics that separated the cell populations across time were enriched for *chromosome segregation* (GO:0007059; padj.: 1.89e-78), *mRNA processing* (GO: 0006397; padj.: 7.65e-47), *regulation of synapse structure or activity* (GO: 0050803; padj.: 1.06e-14), and *actin filament organisation* (GO: 0007015; padj.: 7.73e-05), among others (**Fig. 1c**, **Table 1**). This analysis revealed neuron-specific functions that were particularly enriched in the pNeuroD1+ population over time. The NeuroD1 promoter provided a “pure” upper-layer neuronal identity, whereas the CAG promoter mainly labelled upper-layer neurons among cells with glial signatures; therefore, certain clusters diverged (**Fig. 1c**). Overall, our results show that isolated cells with a specific neuronal promoter differ in the transcriptome landscape compared with cells, which were isolated using a general promoter (**Supp. Fig. 1g**). These results emphasise the relevance of our cell-specific approach in delineating the molecular background of a defined neuronal population differentiating in the cortex.

We therefore focused our analyses on the cell population isolated using promoters specific to neuronal progenitors (pGlast1+) and early developing neurons (pNeuroD1+). Clustering was designed to identify genes whose expression dynamics changed over time within the specific cell population, an effect that was obscured when both populations were analysed together. This analysis of pGlast1+ and pNeuroD1+ cells identified 10,114 DEGs between E14 and P7 (padj < 0.01; **Fig. 2a**, **Table 1**), which were grouped into six major clusters representing gradual transcriptional transitions from progenitors at E14 (pGlast1) through early developing neurons at E16-E19 (pNeuroD1) to mature ULNs at P3, P5, and P7. These clusters capture the principal molecular programmes active during this developmental window. For example, GO terms such as “*RNA splicing”* (Cluster 1, GO:0008380; padj.: 1.29e-78) are upregulated prenatally but become downregulated after birth, including *Prmt1* and neuronal progenitor marker genes such as *Neurog1* and *Eomes*. The expression of genes associated with axonogenesis and neuron projection guidance peaks between E17 and E18 encompassing genes like the upper-layer marker *Satb2* (Cluster 2, **Fig. 2a**). Cell cycle genes such as *Ccnb1* fall below average gene expression after E16 (Cluster 3, **Fig. 2a**). In contrast, GO terms such as “*regulation of synapse structure or activity”* with genes such as *Grin2b* (Cluster 4, GO:0050803; padj.: 1.18e-14) show the opposite pattern (**Fig. 2a**). Chemotaxis and catabolic processes, including *C1qa* and *Wnt7a* representative expression, are subsequently upregulated towards mature stages (Clusters 5 and 6, **Fig. 2a**). At birth (E19_E, prenatal; E19_P, postnatal), most of the trajectories of the analysed gene clusters converged and subsequently underwent a marked shift in expression direction after birth (**Fig. 2a, Supp. Fig. 2a**), suggesting a critical developmental transition in the molecular landscape.

**Figure 2.**
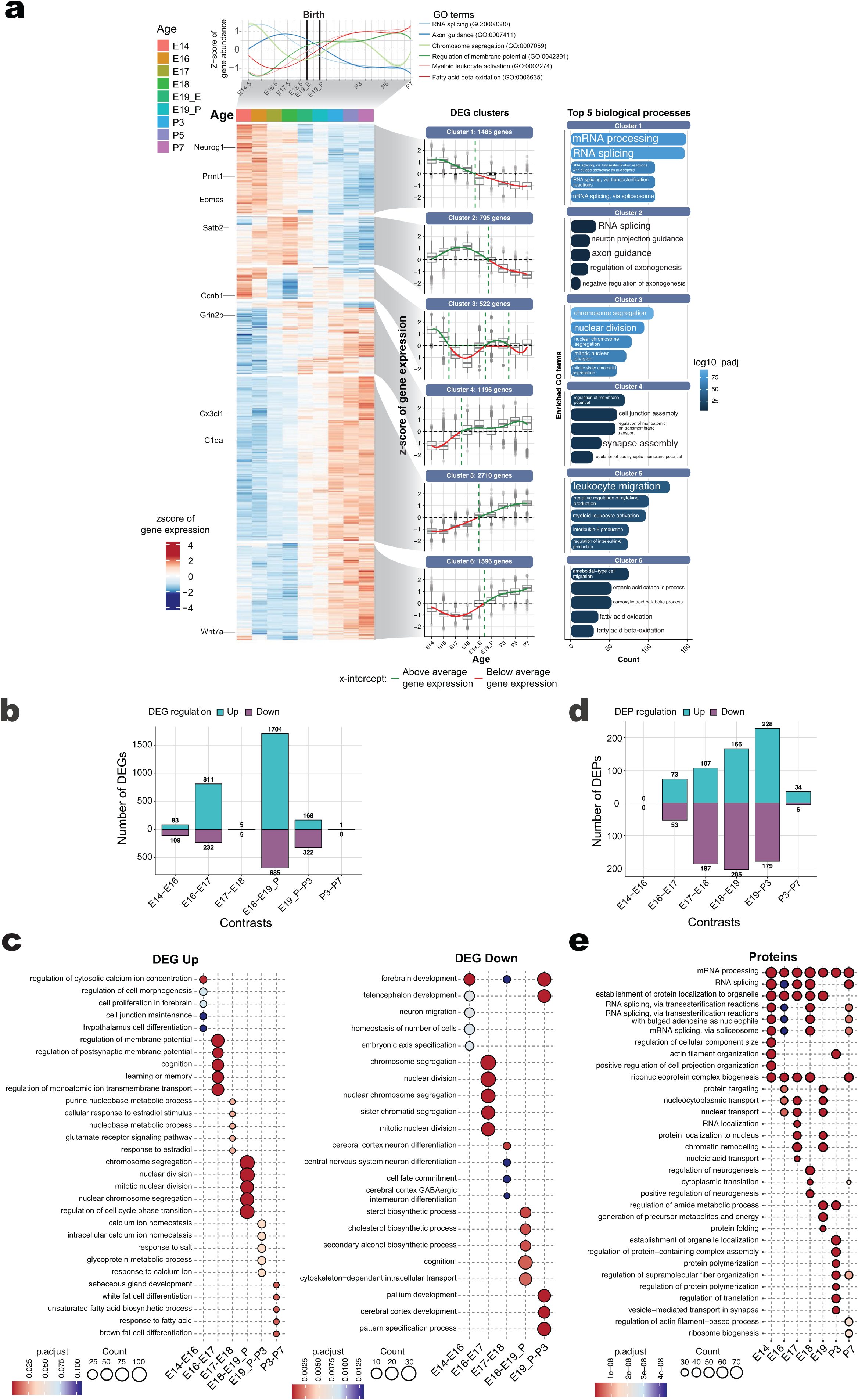
Birth-associated molecular transition in the pGlast1⁺/pNeuroD1⁺ population. **a**) *Temporal dynamics of the upper-layer neuronal population indicate six major gradual shifts during development.* By restricting analysis to the pGlast1+/pNeuroD1+ population, six expression clusters capture a continuous progression from neuronal progenitors (E14) to mature neurons (P7). Z-scored gene expression trajectories reveal a coordinated shift in most clusters from below-to above-average expression around birth (E19), indicating a major transcriptional transition. RNA splicing-associated genes are progressively downregulated, whereas catabolic processes are upregulated. Axonogenesis peaks at E18, nuclear division is cyclic, but declines after E16, and genes linked to membrane potential regulation, synapse assembly, and leukocyte migration are upregulated postnatally. **b**) *The transcriptome does not change during migration (E17-E18) and postnatally*. Consecutive DEG comparisons from E14 to P7 of the pGlast1+/pNeuroD1+ population show a transcriptional burst during early neurogenic transitions (E14-E16) and at birth (E19), whereas few changes are detected during migration or after postnatal day 3 (P3). **c**) *Associated GO terms outline the gradual acquisition of neuronal functions and the loss of progenitor character*. Early transitions (E14-E17) are characterised by upregulation of morphogenesis and homeostatic programmes, followed by enrichment of neuronal excitability and cell cycle regulation mid-embryonically, and metabolic specialisation postnatally (top five GO:BP terms per transition). Downregulated genes are predominantly associated with early neurodevelopmental patterning, forebrain development, neuronal differentiation, and cell fate commitment, consistent with progressive shutdown of developmental programmes. **d**) *The proteome accumulated changes until birth, after which it does not change further.* Consecutive proteomic comparisons across developmental stages reveal an increase in differentially detected proteins approaching birth, followed by a postnatal period of absence of detectable proteomic changes. **e**) *During cell maturation, RNA splicing progressively decreases at the protein level, as the cell specialises in postsynaptic protein regulation.* The upper-layer neuronal proteome was stratified by stage-specific relative abundance, revealing increased postsynaptic organisation and reduced RNA splicing processes towards P7 (top five GO:BP terms per stage).

Notably, the only synchronised upregulation during this perinatal window was associated with cilium organisation and function (**Supp. Fig. 2b**). Cilium-related GO terms (213 associated genes; e.g. GO:0044782, “*cilium organisation”*) exhibited a pronounced local maximum in expression around birth. During the E18-P3 interval, “*cilium movement”* emerged as the most highly enriched GO term, whereas *RNA splicing* was the least enriched; “*synaptic processes”* were also significantly enriched, albeit to a lesser extent (GSEA, **Supp. Fig. 2b**). When the entire developmental window from E14 to P7 was considered and full GO terms (i.e. all associated genes) were analysed, cilia-associated GO terms – including “*cilium organisation”* – again displayed a local maximum, as shown by GSVA (**Supp. Fig. 2b**).

GO analysis of DEGs across successive developmental transitions revealed distinct, stage-specific biological programmes that are overrepresented during upper-layer neurogenesis. Genes upregulated during early embryonic stages (E14-E17) were predominantly enriched for processes related to cell morphogenesis, extracellular matrix organisation, and ion homeostasis. In subsequent mid-embryonic transitions (E16-E17 and E18-E19_P), enrichment shifted toward pathways regulating neuronal excitability, membrane potential, and cell cycle progression. Postnatally (P3-P7), upregulated genes were primarily associated with lipid metabolic processes. In contrast, downregulated genes showed progressive enrichment for early developmental programmes, including forebrain and telencephalon development, neuronal differentiation, cell fate commitment, and mitotic cell cycle regulation, indicating their systematic repression as development proceeds (**Fig. 2b, c**).

At the proteomic level, we observed similar gradual shifts in protein abundance from E16 to P3 (**Fig. 2d**). Prenatally, these proteins are predominantly associated with RNA splicing. Our results reinforce the notion that RNA splicing is a central cellular process underpinning early neuronal differentiation^19–23^. In contrast, at birth and postnatally, they relate mainly to synapse organisation and function (**Fig. 2e**).

Until birth, the global transcriptome of developing neurons does not substantially diverge from their progenitor state (pGlast+ cells at E14, **Supp. Fig. 1g**). Early-developing neurons (E16-E18) retained transcriptional profiles closely resembling those of E14 progenitors. Thus, despite exhibiting morphological hallmarks of neuronal differentiation – such as migration and axon formation (**Fig. 1a**) – these neurons maintain a progenitor-like molecular identity.

In summary, DEG analyses indicate that upper-layer neurogenesis involves a coordinated progression of molecular events, including transcription factor networks and post-transcriptional and post-translational maturation of mRNA and proteins.

To identify shared and discordant molecular programs between the mRNA and protein layers across developmental time, we performed an integrative analysis of our matched transcriptomic and proteomic datasets (**Fig. 3a**). We used a supervised multiblock Partial Least Squares Discriminant Analysis (PLS-DA) focused on the pGlast+/pNeuroD+ cell population, followed by loading-based feature selection and stage-wise trajectory modelling. As mentioned earlier, the RNA-seq dataset, along with the supervising “Age”/developmental stage factor, provided the necessary scaffold for the protein dataset to extract key concordantly regulated molecular pathways (**Fig. 3** and **Supp. Fig. 1f**).

**Figure 3.**
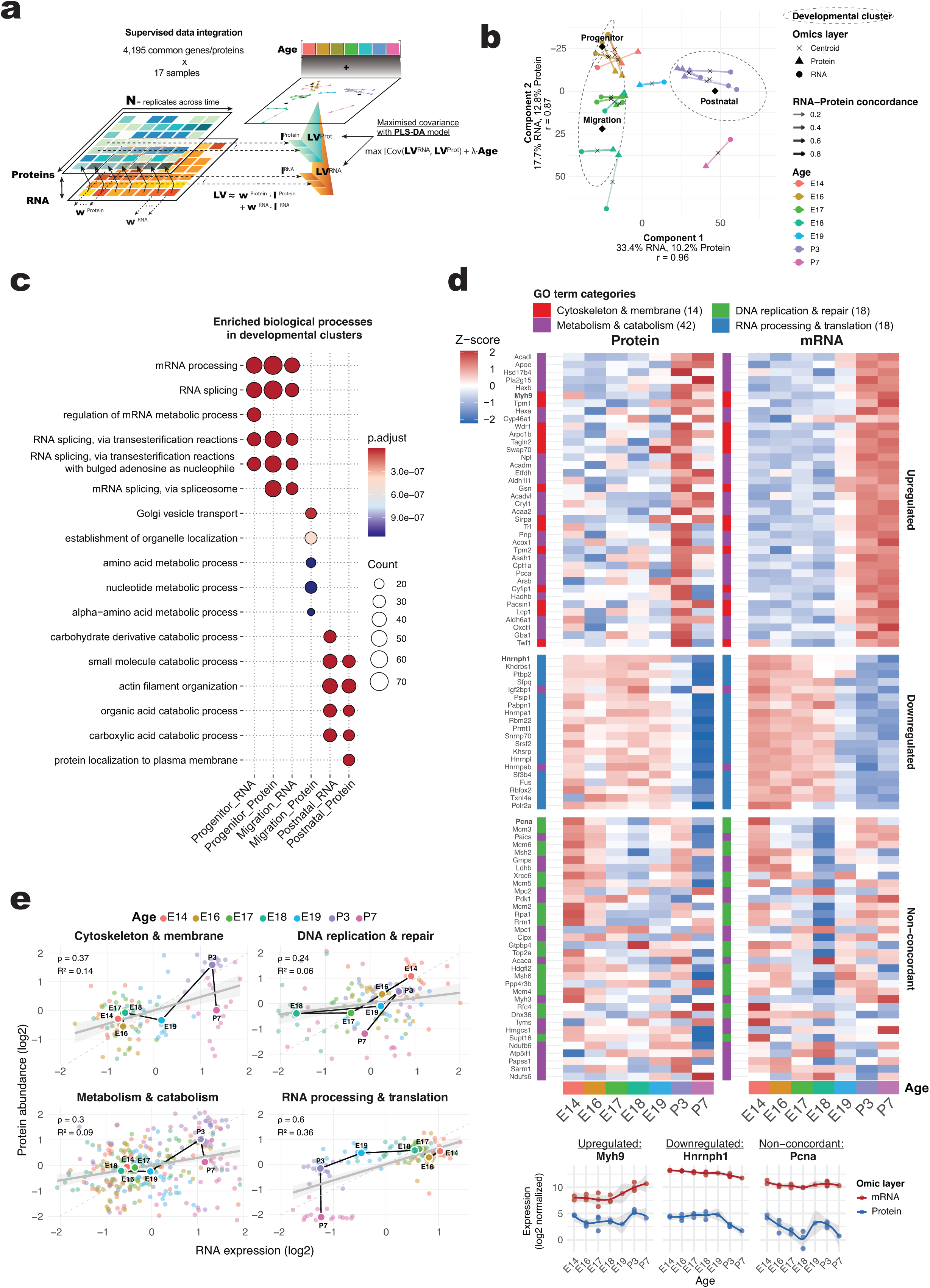
Integrated transcriptomic–proteomic analysis reveals coordinated repression of RNA splicing and activation of extracellular matrix reorganisation. **a)** *Cross-omic integration strategy.* RNA-seq and matched protein abundance data (4,195 shared features across 17 samples, E14-P7) were integrated using block PLS-DA (mixOmics). RNA and protein data were treated as separate layers and jointly decomposed into latent components, maximising covariance between these omic layers while supervising component extraction by developmental stage. For each component, sample scores per omic layer were obtained by projecting each omic layer onto its respective loading vectors. The resulting latent variables define the corresponding latent variable spaces (LV^RNA^, LV^Protein^), which summarise the dominant sources of variation within each omics layer. These latent variables are jointly estimated to maximise covariance between the RNA and protein layers, weighted by the design parameter λ (λ = 0.1), while developmental stage is included as a categorical outcome to supervise component extraction towards stage-discriminative variation. This way, the contribution of individual genes or proteins to the components was defined. **b)** *Concordant RNA-protein trajectories across development.* Projection onto the first two PLS-DA components shows stage-dependent progression. Samples correlate between mRNA and protein layer (RNA-protein variation for component 1: r = 0.96) and cluster into progenitor (E14-E16), migration (E17-E18), and postnatal (E19-P7) phases, with centroids and confidence ellipses highlighting coordinated developmental trajectories. **c)** *Stage-specific biological programmes conserved across omic layers.* Correlated biological processes between mRNA and protein layers are enriched in RNA splicing, which predominates at progenitor stages, in Golgi vesicle transport during migration, and postnatally in small-molecule catabolism, actin filament organisation, and extracellular matrix remodelling. This stage-dependent transcriptional and proteomic regulation appears tightly coupled during neuronal maturation. **d)** *Concordant and discordant RNA-protein dynamics.* Heatmaps show z-scored RNA and protein abundance across E14-P7 upper-layer neurogenesis, highlighting representative concordant features. Major GO biological process categories for upregulated, downregulated, and non-concordant genes and proteins are annotated, with the number of transcripts or proteins in each category indicated in brackets. Genes and respective proteins are plotted below across age and omic layer. **e)** *Functional stratification of RNA-protein concordance.* RNA-protein relationships were summarised across GO:BP categories (Cytoskeleton & membrane; RNA processing & translation; DNA replication & repair; Metabolism & catabolism). Stage-wise trajectories reveal the strongest concordance for cytoskeletal and metabolic genes, and the weakest concordance for DNA replication and repair. Dashed line indicates theoretical perfect concordance; correlations are quantified by Spearman’s ρ across developmental ages, with indicated developmental stage-wise medians connected with a line.

We identified two major latent components that captured most of the shared variation across developmental stages (**Fig. 3b**). Their component scores revealed temporal clusters of samples corresponding to progenitor (E14-E16), migration (E17-E18), and postnatal stages (E19-P7) (**Fig. 3b**, **Table 2**).

Progenitor-stage genes, which declined towards P7, were strongly enriched for RNA processing and splicing at both RNA and protein levels. Key GO terms included “mRNA processing” (GO:0006397, RNA padj = 6.19e-25; protein padj = 1.27e-46), “RNA splicing” (GO:0008380, RNA padj = 1.94e-22; protein padj = 3.26e-48), and more specific spliceosome-related processes such as “RNA splicing, via transesterification reactions” (GO:0000375/0000377, RNA padj = 1.12e-17; protein padj = 3.67e-40) and “mRNA splicing, via spliceosome” (GO:0000398, RNA padj = 1.12e-17; protein padj = 3.67e-40). Regulatory processes, including “regulation of mRNA metabolic process” (GO:1903311, RNA padj = 2.30e-19; protein padj = 3.38e-26), “regulation of RNA splicing” (GO:0043484, RNA padj = 1.29e-14; protein padj = 4.54e-25), and “chromatin remodelling” (GO:0006338, RNA padj = 2.12e-15), were also strongly represented, reflecting active transcriptional and post-transcriptional control. Additional protein-level enrichment included “ribonucleoprotein complex biogenesis” (GO:0022613, padj = 1.50e-25) and “RNA localisation” (GO:0006403, padj = 3.28e-20).

Migration-stage genes displayed heterogeneous RNA-protein dynamics, particularly at E18. RNA-level enrichment remained focused on RNA processing and splicing, including “mRNA processing” (GO:0006397, padj = 1.41e-17), “RNA splicing” (GO:0008380, padj = 1.05e-12), and related spliceosome activities (GO:0000375/0000377/0000398, padj ≤ 4.38e-11), as well as “RNA localisation” (GO:0006403, padj = 4.58e-10) and “regulation of chromosome organisation” (GO:0033044, padj = 1.01e-10). Synaptic and vesicle-related processes were also enriched, including “vesicle-mediated transport in synapse” (GO:0099003, padj = 6.86e-10) and “synaptic vesicle cycle” (GO:0099504, padj = 1.16e-9). At the protein level, migration-stage genes were enriched for “Golgi vesicle transport” (GO:0048193, padj = 2.96e-8), “establishment of organelle localisation” (GO:0051656, padj = 4.20e-7), and multiple metabolic processes, including “amino acid” (GO:1901605), “nucleotide” (GO:0009117), and “pyruvate metabolic processes” (GO:0006090, padj ≤ 2.85e-6), highlighting concurrent cytoskeletal, transcriptional, and metabolic remodelling.

Postnatal-stage genes increased in expression and were enriched for catabolic and metabolic pathways. RNA-level terms included “small molecule catabolic process” (GO:0044282, padj = 3.86e-19), “carbohydrate derivative catabolic process” (GO:1901136, padj = 1.21e-19), “actin filament organisation” (GO:0007015, padj = 3.86e-19), and “fatty acid beta-oxidation” (GO:0006635, padj = 9.25e-11), as well as broader lipid and organic acid catabolic processes. At the protein level, enrichment included “actin filament organisation” (GO:0007015, padj = 6.18e-15), “small molecule and organic acid catabolic processes” (GO:0044282/GO:0016054/GO:0046395, padj ≤ 4.01e-11), “vesicle-mediated transport in synapse” (GO:0099003, padj = 2.59e-10), and processes related to protein localisation and cellular structure, such as “protein localisation to plasma membrane” (GO:0072659, padj = 2.30e-10) and “regulation of actin filament-based process” (GO:0032970, padj = 2.41e-10), reflecting the transition to postnatal differentiation and metabolic adaptation (**Fig. 3c**).

Linear regression of these gene/protein trajectories from E14 to P7 classified 421 concordant genes/proteins into three patterns: monotonically upregulated (n = 125), monotonically downregulated (n = 87), and non-monotonic or non-concordant (n = 209), ranked by their maximal absolute loadings. These genes were identified through component loading analysis, which revealed a limited subset with strong contributions to the shared latent structure. Applying an 80th-percentile threshold to absolute loadings in both the RNA and protein layers resulted in genes/proteins that contributed consistently across components and exhibited coordinated RNA-protein behaviour, indicative of robust multi-omic regulation during development.

Upregulated genes that also increased their protein output across development include representative examples involved in motor protein regulation of cytoskeletal dynamics, migration, and cell adhesion dynamics. For example, the GO terms “small molecule catabolic process” (GO:0044282, padj = 1.94 × 10⁻¹³), “actin filament organisation” (GO:0007015, padj = 3.53 × 10⁻), and “response to extracellular stimulus” (GO:0009991, padj = 3.68 × 10⁻) are associated with Myh9 as a representative gene/protein.

Highlighting suppression of RNA metabolic processes towards birth (**Fig. 3d**), the consistently downregulated cluster at mRNA and protein level encompassing, e.g. *Hnrnph1*, was enriched for *“RNA splicing”* (GO:0008380, padj = 2.60 × 10⁻¹¹), *“mRNA processing”* (GO:0006397, padj = 8.14 × 10⁻), and related spliceosome-mediated functions.

Non-concordant genes exhibited heterogeneous, uncoupled RNA-protein dynamics despite high loadings, such as *Pcna*, and were primarily associated with *“DNA replication”* (GO:0006260, padj = 2.57 × 10⁻) (**Fig. 3d**). Given their strong contribution to the shared latent structure at either the RNA or protein level, the non-concordance may indicate regulation beyond transcription.

Functional enrichment across these three patterns of RNA-protein concordance corresponded to RNA processing and translation (downregulated; ρ of mRNA-protein correlation ≈ 0.60), cytoskeleton and membrane organisation (upregulated; ρ ≈ 0.47), DNA replication and repair (non-concordant; ρ ≈ 0.24), and metabolism and catabolism as a heterogeneously regulated process (ρ ≈ 0.29) (**Fig. 3e**). Together, this corresponds to what can be derived from the modalities when analysed individually, so PLS-DA effectively identified the subset of transcripts and proteins that jointly encode developmental progression.

Altogether, the concordant genes and proteins show a coherent, synergistic multi-layer biological programme, resulting in a rapid response during this developmental phase from mRNA transcript to the implementation of protein expression. Building on RNA-seq and proteomics analyses, we observed gradual transitions from progenitor to mature upper-layer neurons between E14 and P7. Early neurons retained a transcriptional profile similar to their E14 state, even as neurons began to exhibit morphological hallmarks of differentiation, while postnatal neurons displayed distinct, stage-specific transcriptional and proteomic programs. Differentially expressed genes clustered into coherent temporal modules, reflecting a shift from RNA processing and splicing in early stages toward cytoskeletal remodelling, synaptic organisation, and metabolic adaptation postnatally. Across all clusters, coordinated RNA-protein dynamics highlight profound transcriptional reprogramming during differentiation^8^, with some processes (e.g., cilium organisation) peaking perinatally. Overall, these results reveal a gradual, multi-layered acquisition of neuronal identity, with progenitor-like molecular signatures persisting in differentiating neurons before converging toward mature functional states.

### Neurogenesis upon Maternal Immune Activation

MIA has been implicated in increasing the risk of neurodevelopmental disorders in offspring^11–14^. Therefore, we decided to test whether MIA alters the molecular landscape of early developing neurons. To investigate how MIA affects the development of ULNs, we subjected them to this condition with PolyI:C and monitored the subsequent molecular changes in relation to our findings in the unperturbed state (**Fig. 4a; Suppl. Fig. 4a**). First, the sera of mother mice subjected to MIA were tested for elevated cytokine levels, as these indicate actual immune activation and can pass the placental barrier^24,25^ (**Suppl. Fig. 4b**). Maternal serum cytokine levels following PolyI:C administration revealed a significant increase in IL2, IL1b, and IL10 compared to controls, showed a robust activation of both pro-inflammatory and regulatory pathways, while IFNg, IL17a, MCP1, and TNFa showed no significant alteration suggesting moderate immune stimulation without systemic inflammation. Accordingly, the experimental mice gained weight consistent with pregnancy in the days following injection (PolyI:C and control conditions, **Suppl. Fig. 4c; Table 3**).

**Figure 4.**
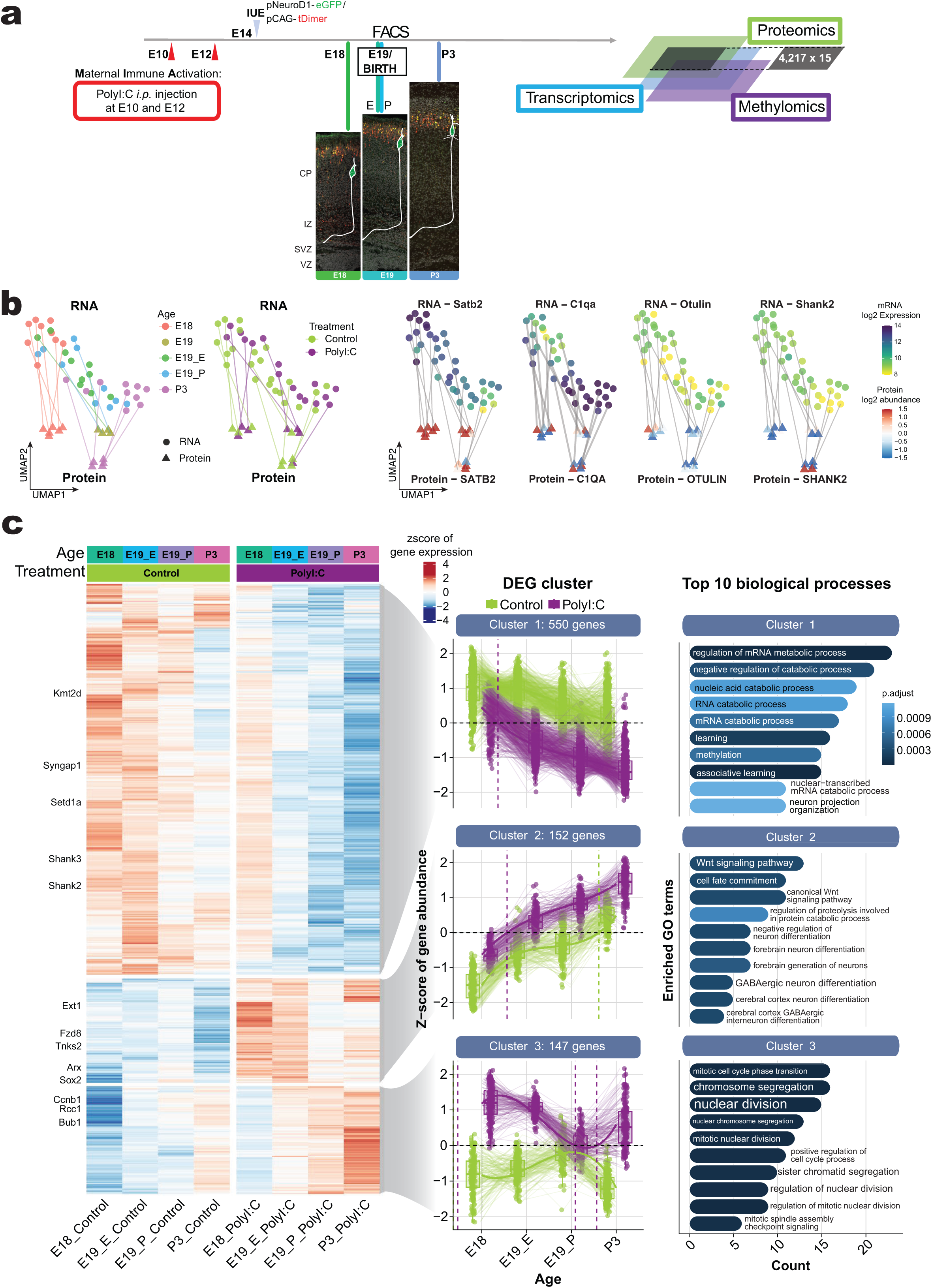
Maternal immune activation induces pathways guided by Wnt signalling at the transcriptomic level. **a)** *Experimental design for maternal immune activation protocol.* Pregnant dams received PolyI:C injections at E10 and E12, followed by *in utero* electroporation and cell sorting congruent to Fig. 1a. Analyses were restricted to pNeuroD-eGFP+ cells at E18, E19, and P3, with DNA methylation additionally assessed at E18. **b)** *Representative transcriptomic and proteomic responses to maternal immune activation.* UMAP embeddings of mRNA expression and protein abundance show developmental trajectories of control and PolyI:C-treated samples at E18, E19, and P3 (proteins: E19 only). Matched mRNA-protein measurements are connected, highlighting divergent responses to maternal immune activation and unchanged responses with respect to the control population. Representative mRNA and protein expression profiles are shown (*Satb2*, *C1qa*, *Otulin*, and *Shank2*), capturing either the Age progression (*Satb2, C1qa*) or differences in treatment condition (*Otulin* vs. *Shank2*). **c)** *PolyI:C-induced gene expression programmes during corticogenesis.* Differential expression analysis identified 849 PolyI:C-responsive genes across development. Clustering reveals three major response patterns: (i) sustained repression of catabolic, learning, and neuron projection organisation genes from E18 onward; (ii) precocious and persistent upregulation of Wnt signalling, forebrain neuron differentiation, and cell fate commitment genes relative to controls; and (iii) prolonged activation of nuclear division and cell cycle regulatory genes. Enriched GO biological processes illustrate treatment-specific deviations from normal developmental trajectories.

To further characterise the effect of MIA, we performed RNA-seq and proteomic analyses of the offspring following the previous pattern. UMAP embeddings of the PCA outputs highlight the 15 matched biological pNeuroD+ samples of both mRNA and protein layer in control and MIA conditions, which are connected by lines to indicate correspondence (**Fig. 4b**). Examples of genes and their respective proteins show time-dependent down– or upregulation upon PolyI:C treatment (**Fig. 4b**).

Cluster analysis of the most significant genes upregulated or downregulated shows that MIA treatment caused the downregulation of key synaptic genes like *Syngap1*, *Shank2,* and *Shank3* (Cluster 1), which are linked to GO terms such as “*learning”* (GO:000761, padj.: 0.000173), “*associative learning”* (GO:0008306, padj.: 3.06e-06, **Supp. Fig. 5a**), “*neuron projection organisation”* (GO:0106027, padj.: 0.001155), “*methylation”* (GO:0032259, padj.: 0.000594), and “*regulation of mRNA metabolic process”* (GO:1903311, padj.: 3.06e-06), among others (**Fig. 4c**; **Supp. Fig. 5b**; **Table 1**). Conversely, we observed upregulation of genes such as *Fzd8*, *Otulin*, *Tnks2*, *Arx*, *Sox2*, and *Ccnb1* (Clusters 2 and 3), which are associated with GO terms such as “*Wnt signalling pathways”* (GO:0016055, padj.: 0.000286), “*canonical Wnt signalling pathway”* (GO:0060070, padj.: 0.000283, **Supp. Fig. 5a**), “*nuclear division”* (GO:0000280, padj.: 5.95e-06), “*chromosome segregation”* (GO:0007059, padj.: 9.30e-07), and “*mitotic cell cycle phase transition”* (GO:0044772, padj.: 9.30e-07), among others (**Fig. 4c**; **Supp. Fig. 5b**). These findings suggest that neurons exposed to MIA may delay neuronal differentiation.

**Figure 5.**
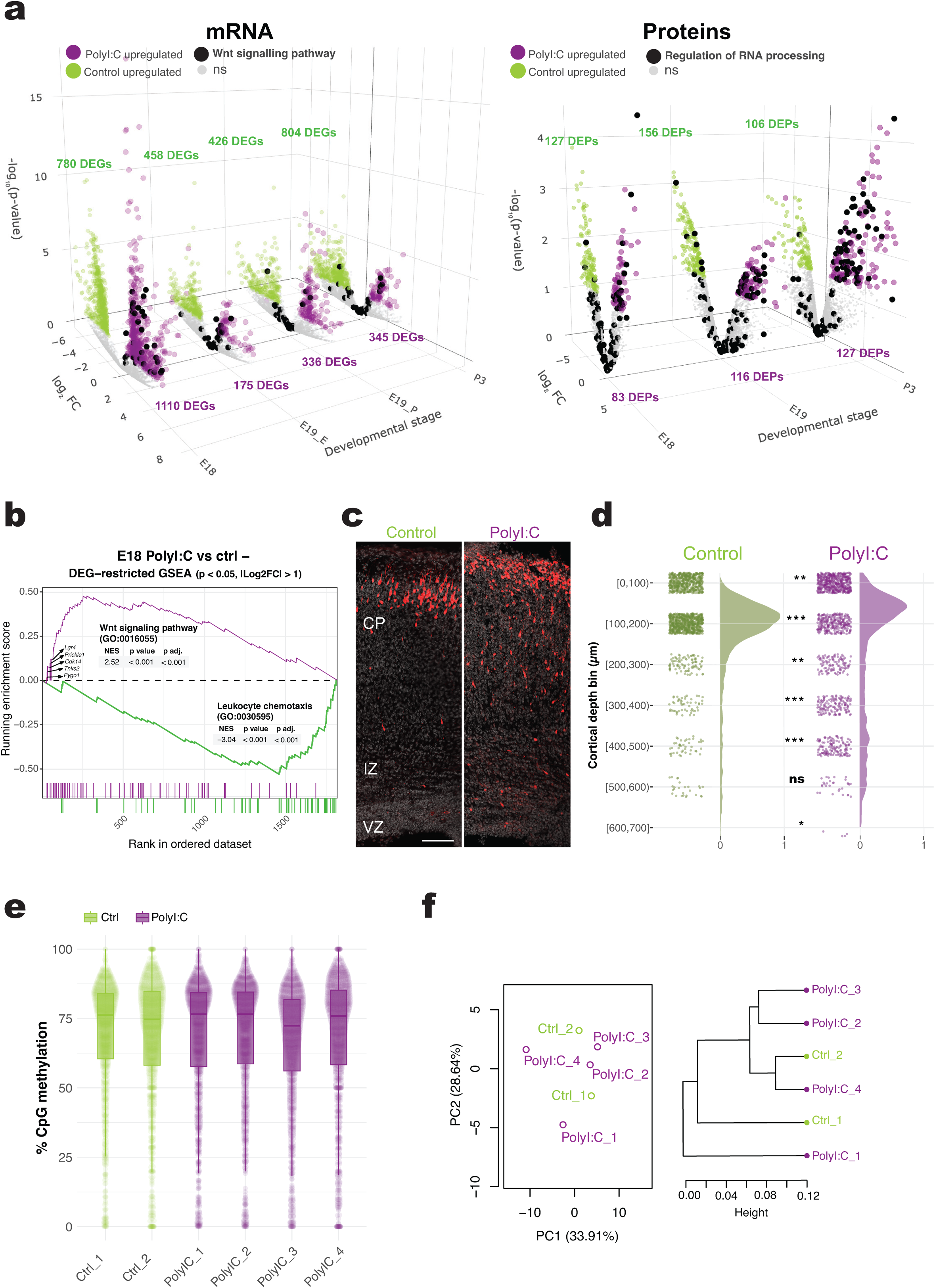
Maternal immune activation induces opposing regulation of canonical Wnt signalling and synaptic translation at transcriptomic and proteomic levels. **a)** *Differential responses to PolyI:C treatment.* Volcano plots highlight significantly regulated mRNAs (left) and proteins (right) in PolyI:C-treated versus control pNeuroD+ cells (adjusted *p* < 0.05, |log₂FC| > 1). Features upregulated by PolyI:C are shown in magenta, and control-enriched features in green. Wnt-associated transcripts and RNA processing proteins are annotated (black). **b)** At E18, the control cohort predominantly upregulates baseline leukocyte chemotaxis pathway genes (as shown in Figure 2a), whereas the PolyI:C cohort exhibits distinct Wnt pathway activation: GSEA of transcriptional changes at E18 reveals significantly enriched pathways (Wnt signalling (GO:0016055) and leukocyte chemotaxis (GO:0016055)) from the PolyI:C versus control DEG comparison (p < 0.05, |log₂FC| > 1) that were ranked using a signed –log₁₀(p-value) metric. **c)** Representative images of tDimer+ cells at E18 in coronal sections from PolyI:C and untreated animals reveal more cells in the PolyI:C condition in the lower cortical plate (CP). At this time, cells should have completed migration from the ventricular zone (VZ) to the CP as seen in the control condition. Scale bar = 100 µm. **d)** *Quantification of cell distribution across cortical depth.* Binned counts (100 µm) show an accumulation of tDimer+ cells in PolyI:C-treated cortices at 300–500 µm bins, suggesting delayed migration compared with controls. Adjacent density plots summarise distributions. Statistical significance per bin was assessed by Fisher’s exact test (*p < 0.05; **p < 0.01; ***p < 0.001; ns = not significant), and overall distributions by GLMM with LRT. **e)** *Per-sample distributions of CpG methylation percentages.* Samples from embryos of PolyI:C-treated mother mice (magenta) show profiles that highly overlap with those of the control group (green). Median methylation remains close to 75% across all six samples, and interquartile ranges largely coincide. The extensive overlap of individual CpG-level values further highlights the absence of a condition-specific methylation signature at the whole-genome level. **f)** *Correlation between global methylation profiles.* Neither PCA shows separation between PolyI:C-treated and control samples, indicating that the major sources of variance in methylation are not condition-dependent (PC1 and PC2 explain the largest proportion of variance), nor clustering using correlation distance and Ward linkage segregate PolyI:C-treated and control samples, indicating high similarity in global methylation profiles across conditions.

Most of the significantly up– or downregulated genes following MIA treatment were identified at E18 (**Fig. 5a**; **Table 4**). However, consistent proteomic alterations were observed across all analysed time points (**Fig. 5a, Supp. Fig. 6a**). Our data show that genes associated with GO terms such as “*Wnt signalling pathway”* and “*canonical Wnt signalling pathway”* were upregulated in response to MIA at E18 (**Fig. 5a, b**; **Supp. Fig. 5b, Supp. Fig. 6b**). In contrast, under control conditions, genes related to immune response were more expressed compared with MIA-treated neurons (**Fig. 5b**). In neurons, many genes that are categorised under GO terms such as *“immune response”* are in fact involved in synaptic pruning and plasticity, as exemplified by the previously mentioned ***C1qa*** (**Fig. 4b**), which is indeed more expressed in the control group than in the PolyI:C group. Likewise, factors annotated under GO terms such as *“leukocyte chemotaxis”* are likely to participate in growth cone repulsion mediated by semaphorins and ephrins during neuronal migration and subsequent integration into pre-existing circuits established by earlier-born neurons^26–30^.

At the proteomic level, we did not detect differential expression of proteins associated with Wnt pathways at E18 or other time points, which may reflect limitations of the mass-spectrometry workflow. Wnt pathway proteins – including secreted Wnt ligands, lipid-modified proteins, and membrane-associated receptors – are notoriously difficult to detect by standard MS-based proteomics due to their hydrophobicity and poor recovery during sample preparation^31,32^.

Notably, proteins involved in RNA splicing were upregulated upon MIA treatment at E18 and P3 (**Fig. 5a**), in contrast to control conditions in which *RNA Processing & Translation* steadily decreased over time (**Fig. 3e**). These findings suggest that MIA induces a sustained shift in post-transcriptional regulatory mechanisms. Although Wnt-related proteomic changes were not detected, the consistent alteration of RNA-splicing proteins highlights a potentially underappreciated layer of molecular dysregulation. This is particularly notable as RNA splicing was the most downregulated group of genes in the unperturbed condition at both the protein and mRNA levels and is probably a key driver of neuronal differentiation (**Fig. 3**).

Comparing the gene expression patterns between E18 and P3 reveals that not all genes maintain consistent trajectories across development. Nevertheless, genes associated with the Wnt pathway remain highly expressed at both E18 and P3 following MIA treatment (**Supp. Fig. 7a, b**). Together, these results point to RNA-processing pathways as candidate mediators of MIA-driven developmental effects and highlight that MIA induces both persistent and stage-specific molecular alterations in the developing cortex. While Wnt-related transcriptional changes appear sustained across late embryonic and early postnatal periods, immune-related signatures shift dramatically over time, suggesting dynamic and context-dependent responses to early inflammatory exposure. Such temporal heterogeneity underscores the importance of considering the developmental stage when assessing the long-term consequences of MIA on cortical maturation.

Next, we examined whether the transcriptomic changes we identified translate into functional effects at the cellular level. Previous studies have shown that precise regulation of the canonical Wnt signalling pathway influences neuronal migration^33^. At E18, we observed an upregulation of several genes associated with canonical Wnt signalling (**Supp. Fig. 6b**), prompting us to investigate whether neuronal migration is altered following MIA exposure. To this end, we performed *in utero* electroporation at E14 to introduce the pCAG-tDimer/pNeuroD-eGFP plasmids and collected brain tissue at E18 for analysis. Our results indicate that the positioning of pCAG-tDimer+ migrating neurons is disrupted in MIA-treated brains compared with controls. Specifically, a greater proportion of labelled cells were located within the IZ and lower CP, compared between MIA-treated and control, consistent with delayed neuronal migration (**Fig. 5c, d**). While the median position of the migrating neurons does not greatly differ between conditions, a quantile-based analysis reveals a great shift in the proportion of cells in the lower CP and IZ (**Supp. Fig. 8**). Altogether, these results indicate that MIA exposed developing brains exhibit neuronal migration deficits driven by alterations in their molecular landscape, probably in particular within Wnt signalling pathways.

Finally, we assessed whether MIA alters the methylome landscape of differentiating ULNs at E18. Global DNA methylation profiles were highly similar between PolyI:C-treated and control brains, with no detectable condition-specific differences (**Fig. 5e, f; Supp. Fig. 9**). Per-sample CpG methylation distributions and tile-based analyses showed comparable patterns across conditions, and the four PolyI:C-treated and two control samples exhibited high overall correlation (r ≈ 0.87-0.98, **Fig. 5e**, **Supp. Fig. 9e**, **Supp. Fig. 9d**), with no separation by condition in unsupervised PCA or clustering analyses (PC1 and PC2 explaining 33.91% and 28.64% of the variance, respectively, **Fig. 5f**). Although CpG coverage varied substantially across samples (PolyI:C: 7,184-16,067 CpGs; controls: 4,165-15,440 CpGs, **Supp. Fig. 9c**), only 1,004 CpGs were shared across all six samples. Among these shared sites, 168 CpGs (16.7%) displayed modest differential methylation (|Δ| > 5%, q < 0.05), with hypomethylation more frequent than hypermethylation. Of the 1,004 CpGs tested, 168 (16.7%) were significantly altered, with hypomethylation predominating (**Supp. Fig. 9f**, 115 hypomethylated vs 53 hypermethylated). Overall, fewer than 0.05% of CpGs were affected, supporting global methylation stability despite maternal PolyI:C treatment. These changes were sparse and failed to cluster into robust differentially methylated regions, and consistent with this, tile-based analysis at 1 kb resolution (|Δ| > 10%, q < 0.1) identified no differentially methylated regions (not shown). This supports the finding that even our candidate genes, such as *Fzd8*, which was upregulated at both the transcriptomic and proteomic levels, did not show differences in methylation coverage between the two conditions (**Supp. Fig. 9g**). Together, these data indicate that MIA does not induce strong global or regional DNA methylation changes in ULNs at E18, although a slight predominance of isolated hypomethylation events suggests subtle chromatin effects. Our multi-omics and functional analyses, however, reveal pronounced transcriptomic alterations – most notably the upregulation of canonical Wnt signalling components – accompanied by defects in neuronal positioning, demonstrating that MIA profoundly reshapes the developmental trajectory of ULNs by disrupting the balance between proliferation, differentiation, and migration during corticogenesis.

## Discussion

The transition of neurons from a proliferative state to terminal differentiation represents one of the most pivotal steps in brain development. Although it is well known that neurons permanently exit the cell cycle upon differentiation, the molecular programs guiding this transition, particularly in upper-layer neurons, remain only partially understood. The study described here addresses this gap by charting the developmental trajectory of murine ULNs using a multi-omics framework.

By integrating transcriptomic and proteomic data, we characterise the evolving molecular landscape of ULNs from their birth to the onset of circuit formation. This approach not only enriches our understanding of normal neuronal maturation but also provides a high-resolution view of the dynamic changes across multiple regulatory levels. Importantly, the multi-omics strategy enables the identification of convergent or divergent patterns across gene expression and protein abundance, offering deeper mechanistic insight than either method alone could provide. Recent studies focused on bioinformatic differentiation between the neocortical layers, which is based on marker genes. We showed that, particularly during developmental processes, these allocations are not always accurate and require a specific purification process as we applied here.

We identified a highly coordinated transcriptome-proteome axis underlying UL neurogenesis. RNA and protein components converge on a shared developmental trajectory (r ≈ 0.96 for LV1) and highlight a compact set of concordant genes with markedly improved cross-omics coherence (421 genes/proteins with 9.1-fold correlation improvement compared to the total input set; see **Supp. Fig. 3**). These results reveal coupled molecular programmes driving embryonic-to-postnatal transitions in upper-layer neuronal maturation.

Based on their relative expression patterns, RNA-splicing – and actin-filament – related processes co-varied across both RNA and protein layers, suggesting that they constitute major molecular drivers of upper-layer neuron (ULN) maturation. In contrast, several cell-cycle-associated features exhibited discordant RNA-protein behaviour, indicating an emerging uncoupling between these molecular layers beginning around the time of neuronal birth. This discrepancy may in part reflect technical limitations, as more differentiated neurons exhibit highly compartmentalised protein synthesis (for example, within dendrites)^34^, which we were unable to isolate. In addition, proteomic resolution remains substantially lower than that of transcriptomics, implying that these cross-layer relationships may be incompletely captured. Consistent with this limitation, the proteomic dataset comprised only a fraction of the treated replicates available for transcriptomic analyses, largely due to the loss of E19 samples during sample preparation.

Because PLS-DA is a linear model, it may not capture the nonlinear, gradual acquisition of cell fate that characterises developmental multi-omics datasets. Although nonlinear approaches could, in principle, model such dynamics more accurately, they require substantially more data points for reliable curve fitting. In our replicate-limited dataset, using the linear coefficients from the PLS-DA model provided a methodologically consistent and robust population-level classification that is less sensitive to sample-specific variation. Our primary interest was to capture directional trends while minimising the risk of overfitting to individual samples.

Biologically, the derived gene groups clustered into coherent functional categories, each comprising genes whose multivariate impact on stage discrimination aligned with characteristic RNA-protein expression matches or mismatches. Each group showed enrichment for one of four principal biological functions. These categories were further supported by stage-specific median RNA-protein expression trajectories. This molecular map of late-born neurons provides a comparative basis for further experimental approaches, such as the MIA approach.

A key strength of this work is its investigation of how environmental perturbations – specifically maternal immune activation (MIA) – influence early neuronal development. As a model of prenatal inflammation, MIA reflects clinically relevant features of maternal infection during pregnancy, a well-established risk factor for neurodevelopmental disorders. The findings show that although the methylome of ULNs remains largely unaffected by MIA, substantial transcriptomic and proteomic changes emerge. MIA induces an upregulation of Wnt signalling pathways alongside a downregulation of key synaptic transcripts. Notably, the absence of alterations in DNA methylation suggests that MIA may disrupt neuronal development primarily through mechanisms affecting translation^35^ rather than gene expression. This dissociation highlights how environmental challenges can shape corticogenesis through epigenetic-independent and post-transcriptional pathways, underscoring the complexity of regulatory processes in early brain development.

One of the most striking observations is the upregulation of Wnt signalling pathways coupled with a downregulation of synaptic developmental programs in MIA-exposed neurons. Wnt pathways are central to early neural patterning, proliferation, and migration, whereas synaptic genes mark progression toward functional circuit integration. The concurrent shift toward amplified Wnt signalling and diminished synaptic transcripts suggests a developmental delay or misspecification of neuronal identity. Such alterations may translate into long-term consequences for cortical connectivity and cognitive function, consistent with the behavioural and structural abnormalities observed in offspring following MIA in other studies^36,37^.

Our study further links these molecular perturbations to disruptions in neuronal positioning during migration – an essential process for establishing proper cortical lamination. However, because our analysis focused only on E18 cortices and did not include earlier stages following the initial PolyI:C administration at E10, we could not determine whether MIA affects neuronal migration directly or instead alters neurogenesis. Nonetheless, this mechanistic connection outlines a plausible pathway through which prenatal inflammation may contribute to neurodevelopmental disorders such as autism or schizophrenia, both of which show epidemiological associations with maternal infection.

In summary, this work offers a significant contribution to our understanding of how early environmental factors shape brain development. By revealing specific signalling pathways and developmental programs disrupted by MIA, the study not only advances basic knowledge of neuronal maturation but also identifies potential molecular targets for early intervention. Future research could build upon these findings by exploring how these initial molecular derangements evolve postnatally and whether therapeutic modulation of pathways like Wnt signalling can restore typical developmental trajectories.

## Acknowledgments

This work was partially supported by a grant PID2024-160995NB-I00 funded by MCIN/AEI/ 10.13039/501100011033, “ERDF/FEDER A way of making Europe” to FCA.

We thank the FACS Core Unit of the University Medical Centre Hamburg-Eppendorf for experimental help and for providing the flow cytometer and sorter used in this study (BD FACSAria Fusion). We thank Robin Scharrenberg for his insightful comments. We thank the Epigenomic, Proliferation and Cell Identity Unit of the Institut Pasteur, Paris for providing the work environment that enabled the completion of the manuscript.

## Methods

### Ethical approval

C57BL/6J mice were reared and housed according to the standard procedures of the Central Animal Facility at the University Medical Centre Hamburg-Eppendorf (UKE), Hamburg. All mouse experiments were conducted in accordance with the German and European Animal Welfare Act. The experimental procedures were approved by the Animal Research Ethics Board, the Hamburg Authority (TVA N051/2020 and N109/2020), and the Animal Welfare Commission of UKE.

All female mice imported from Charles River were allowed to adapt for at least six weeks to avoid transcriptional and epigenetic heterogeneity resulting from different housing conditions^38^. All mice received a standard diet with Altromin (#1311P) ad libitum. After the adaptation phase, female mice were bred in-house with male C57BL/6J mice up to one year old. After overnight mating, the presence of a vaginal plug marked embryonic day 0.5 (E0.5). For clarity, the “.5” has been omitted from all IUE and harvest dates in the manuscript.

### *In utero* electroporations

*In utero* electroporations (IUEs) were performed as described^7^. Four different constructs were used to target distinct cell populations in the developing cortex. For RGC-specific expression^39^, the promoter of the ‘glutamate-aspartate transporter 1’ (Glast1, hereafter also referred to as pGlast) was used with a Kozak sequence to drive expression of the red fluorophore dsRed2 via IUE at E12. The vector had a total size of 5,867 bp (Addgene #17706).

For expression specifically in developing neurons^40^, the promoter of the bHLH transcription factor ‘neuronal differentiation 1’ (NeuroD1, hereafter also pNeuroD) with an adjacent IRES sequence was used to drive expression of eGFP via IUE at E14 (Fig. 4C). The total vector size was 6,911 bp (Addgene #61403).

To target a mixed population, a plasmid containing the non-specific CAG promoter was used. The CAG promoter (pCAG) is a strong, constitutive promoter used in mammalian expression vectors to drive ubiquitous transgene expression^41^. When the construct was introduced via IUE at E12, Venus, a fusion protein of eGFP and eYFP, was expressed under pCAG with a total vector size of 5,454 bp (Addgene #127346). When the construct was introduced via IUE at E14, pCAG drove expression of the red fluorescent protein tDimer, with a total vector size of 7,012 bp (#087 pAAV-CAG-tDimer). The optimal concentration of injected plasmids with developmentally active promoters was 4 μg/μL, whereas for plasmids carrying the pCAG promoter, a concentration of 0.1 μg/μL was sufficient.

## Maternal immune activation

### Injections

Timed-pregnant mice were intraperitoneally (*i.p.*) injected with a final administered concentration of 4 mg/kg PolyI:C (Polyinosinic-polycytidylic Acid Potass., Sigma, P9582-5mg; stock solution 0.05 mg/mL in 1xD-PBS) to induce maternal immune activation (MIA). Repeated stimulations at E10 and E12 were performed to enhance the immune response in the mother mice at the onset of neurogenesis.

The administered PolyI:C concentration was determined to be sublethal (>99% survival rate) in previous experiments by Jacobsen et al., 2021^42^, and was calculated according to the respective weight of each mouse to a final injection volume of less than 200 μL *i.p*. Untreated mice served as controls. Both PolyI:C and control animals were handled primarily in parallel cohorts. Supplementary Figure 4 provides detailed information on the maternal weight of both cohorts. The humane endpoint specified in TVA N109/2020 was defined as a weight loss of more than 25% of the average weight.

All omics experiments were performed in parallel with the untreated samples. For MIA, time points E18, E19, and P3 were selected for the transcriptome and proteome experiments, while E18 was additionally subjected to DNA methylome experiments. Both pNeuroD+ and pCAG+ cell populations underwent identical experimental and sequencing procedures; however, downstream NGS analyses were limited to pNeuroD+ cells.

### Cytokine measurements

To test cytokine levels in the sera of PolyI:C-treated and untreated/control mother mice, a custom Bio-Plex Pro Mouse Cytokine multiplex assay (Bio-Rad, ELISA) was designed and measured using a Bio-Plex 200 system according to the manufacturer’s instructions. The selected cytokines were interferon-γ (IFN-γ), interleukin-1β (IL-1β), interleukin-2 (IL-2), interleukin-6 (IL-6), interleukin-10 (IL-10), interleukin-17α (IL-17α), monocyte chemoattractant protein-1 (MCP-1), and tumour necrosis factor α (TNF-α), mostly consistent with Jacobsen et al., 2021^42^. Four untreated mice and six PolyI:C-treated mice were randomly selected, and their sera, derived from the inferior vena cava, were tested in this assay.

No technical replicates were included. Values were normalised using the standard curve generated for each measured cytokine with the Bio-Plex Manager software. Several values were lost due to underrepresentation (Out of Range Below) of fluorescence as calculated by the Bio-Plex Manager software; these were treated as not available (NA). In contrast, extrapolated values generated by the software were included in the analysis, representing estimated concentrations outside the range of the standards but still within the limits of the fitted eight-point logistic standard curve (**Supp. Fig. 4b**).

### Flow cytometry experiments

A detailed version of the applied flow cytometry protocol can be found in the^7^.

### Proteome analysis

The following preparation of frozen cell aliquots containing 10,000–50,000 cells for mass spectrometry (MS) with label-free quantification (LFQ). The applied settings and parameters for sample preparation are described in^7^.

The primary sequences of the detected peptides were searched using the SEQUEST algorithm integrated into Proteome Discoverer software (v. 3.0.0.757, Thermo Fisher Scientific Inc.) against the murine Swissprot database (January 2023) containing 17,013 entries. As peptide modifications (e.g. post-translational modifications) alter peptide mass, options to consider them in the resulting.msf file were set as follows: carbamidomethylation was set as an artificially induced fixed modification for cysteine residues, while oxidation of methionine, pyroglutamate formation on glutamine residues at the N-terminus of the peptide, and acetylation of the N-terminus of the protein were allowed as biologically induced variable modifications. A maximum of two missed tryptic cleavages was permitted. Only peptides between six and 144 amino acids were considered for further analysis. A strict cut-off (FDR < 0.01) was applied for both peptide and protein identification. Quantification was performed using the Minora algorithm implemented in Proteome Discoverer v. 3.0. The resulting protein abundances were log2-transformed, followed by normalisation of the column means.

Using this bead-based quantitative approach, 4,295 proteins were identified across 74 samples, with consistent protein coverage across cell populations and developmental stages, indicating no significant differences in the number of identified proteins (**Supp. Fig. 3b**). After quality control, six samples were excluded, resulting in 66 samples for downstream analyses.

As the samples were produced in three different batches, batch effect correction was performed using HarmonizR as described by Voss et al. 2022^43^. Thus, further filtering or normalisation steps were unnecessary. However, as the missing values were biased towards less abundant proteins, indicating missing not at random (MNAR) values, the data were imputed using a shift of 0.1 and a scale of 1 (**Supp. Fig. 3c**), which is a subtle imputation to account for MNARs and to avoid artificially inflating differences.

Datasets were stratified by experimental context into WOMIA (no immune activation; 35 samples including the pCAG cell population) and MIA (PolyI:C with age-matched controls; 33 samples including the pCAG cell population), and further subset to pNeuroD+ populations where indicated.

Differential protein abundance was tested across developmental stages (WOMIA: E14-P7; MIA: E18-P3) or between PolyI:C and control within matched ages, using likelihood-based contrasts via the R package DEP (v. 1.22.0; doi: 10.18129/B9.bioc.DEP) with a limma-based algorithm (bayes statistics) and an FDR-based rejection criterion (α = 0.5). Consecutive temporal patterns in Figure 2 are identified by pairwise DEP Wald tests, using a Benjamini-Hochberg adjusted p-value threshold of <0.5 and a log2 fold change either below or above 0.

Given the limited depth and greater variability of the proteomics data, permissive FDR thresholds were applied to capture broad developmental trends rather than strict significance.

### Transcriptome analysis

The transcriptome of frozen aliquots from an average of 10,000 flow-cytometry sorted cells was analysed by RNA-seq.

Total RNA was extracted using the RNeasy Micro Kit (Qiagen, #74004). Before library preparation, the integrity of the extracted total RNA was assessed on an Agilent 2100 Bioanalyzer using a Eukaryote Total RNA Pico assay (Agilent Technologies, #5067-1513). RNA samples with a RIN value below 7 were excluded; however, on two occasions, samples with a RIN below 7 were still approved due to a sufficient level of RNA integrity for sequencing.

For library generation, cDNA was first synthesised, polyA-enriched, and amplified using the SMART-Seq v4 Ultra Low Input RNA Kit for Sequencing (Clontech Laboratories, #634891) according to the manufacturer’s recommendations. The generated cDNA was then further processed using the Nextera XT DNA Library Preparation Kit (Illumina, #FC-131-1096). The fragment length distribution of each library was determined on an Agilent 2100 Bioanalyzer using the DNA High Sensitivity Chip (Agilent Technologies, #5067-4626). All samples were diluted to a concentration of 2 nM, and an equimolar pool of the samples was generated.

1 × 75 bp single-end (SE) sequencing of the samples was performed on an Illumina NextSeq 550 platform, aiming for 20 million reads per sample. A total of 137 samples were bulk RNA sequenced in four runs. The quality of the reads was verified using FastQC (v. 0.11.8; Andrews et al., 2019. http://www.bioinformatics.babraham.ac.uk/projects/fastqc/; last accessed: 09 March 2026). All measured samples passed quality criteria. Nextera adapter clipping was performed using TrimGalore! with cutadapt (v. 3.1; 10.14806/ej.17.1.200). Read mapping was performed using the RNA STAR aligner (v. 2.7.8a;^44^) against the mm10 primary assembly with the corresponding gtf mm10_RefSeq_exon annotation file from the UCSC Genome Browser^45^, using default options for SE raw data. Gap-aware read counting was performed using featureCounts (v. 2.0.1^46^), considering only non-stranded reads that were within an exon and had a minimum MAPQ of 12.

Differential expression analyses were performed using DESeq2^47^ with regularised log-transformed expression (rlog) values.

Overrepresentation analyses were conducted using the clusterProfiler package^48^ with the compareCluster function on Gene Ontology Biological Processes (GO: BP), with genes annotated using the BioConductor data base org.Mm.eg.db (Release 3.22, doi: 10.18129/B9.bioc.org.Mm.eg.db). P values were adjusted using the Benjamini-Hochberg method, and all GO terms were retained (p– and q-value cut-offs set to 1) unless stated otherwise for downstream filtering and interpretation.

Pairwise Wald tests were conducted between consecutive developmental stages, and genes with an adjusted p-value < 0.01 were considered significantly differentially expressed.

Developmental gene expression trajectories were profiled using the R package DEGreport (v. 1.46.0; Pantano, 2025, doi: 10.18129/B9.bioc.DEGreport, https://bioconductor.org/packages/DEGreport; last accessed: 05 May 2026) with the degPatterns function, applied to rlog-transformed expression values of significantly age-associated genes (tested by likelihood ratio test, LRT), which cluster genes by shared temporal patterns across developmental stages.

Differential expression between the cell populations across development was assessed using rlog-transformed counts with an LRT, comparing a full model including age and cell population effects (∼ Age + Cell population) to a reduced age-only model (∼ Age). Genes with adjusted p-values < 0.01 were considered significantly developmentally regulated (n = 5,489). The top 5,000 genes ranked by adjusted p-value were clustered using degPatterns, considering a minimum cluster size of 50 genes.

For visualisation in **Figure 1c**, the vertical dashed lines show the x-intercepts of the respective smoothed profiles, calculated by a fitted polynomial of fifth order to identify gene expression below or above average values. The adjacent bar plots show the top five biological processes (GO: BP) associated with the described DEG clusters, with adjusted p-values indicating the significance of enrichment and the number of considered DEGs for this over-representation analysis.

Developmentally regulated genes focusing on the pGlast+/pNeuroD1+ were identified using an LRT that compares a full model including all covariates to a reduced intercept-only model (∼1). Genes with adjusted p-values < 0.01 were considered significant (n = 10,114). The top 10,000 genes ranked by adjusted p-value were retained for temporal pattern discovery and clustered using DEGreports::degPatterns with a minimum cluster size of 100 genes.

For the MIA condition, count data were analysed with DESeq2, fitting a generalised linear model to only the pNeuroD+ cell population, including age (here focused to E18, E19_E, E19_P, and P3) and treatment (PolyI:C vs ctrl) effects. LRT comparing the full model (design: ∼Age + Treatment) to a reduced model (∼Age) identified 1,010 PolyI:C treatment-associated genes (FDR < 0.05), which were clustered with DEGreport::degPatterns over the Age factor using rlog-transformed expression profiles and considering a minimum cluster size of 50 genes.

Gene set dynamics were analysed using only samples from the pGlast+/pNeuroD+ cell population spanning developmental stages E14 to P7.

Gene Set Enrichment Analyses (GSEA) were performed on this ranked list using the clusterProfiler R package. Mus musculus Gene Ontology Biological Process (GO: BP) gene sets were sourced from MSigDB (v. 2023.2) via the msigdbr R package (v. 25.1.1; Dolgalev, 2025, doi.org/10.32614/CRAN.package.msigdbr, https://CRAN.R-project.org/package=msigdbr; last accessed: 05 May 2026).

Gene set enrichment analysis (GSEA) was conducted using a pre-ranked approach implemented in the clusterProfiler R package. Differentially expressed genes (adjusted p < 0.05, |log₂ fold change| > 1) were ranked by sign(log2FoldChange) × – log10(pvalue). Enrichment was assessed using the Gene Ontology (GO) knowledgebase of gene set-related biological process (GO:BP). Normalised enrichment scores (NES) and multiple-testing-corrected p-values were calculated using the Benjamini-Hochberg method.

Gene Set Variation Analysis (GSVA) was performed with the GSVA package within the Bioconductor framework on rlog-transformed expression values, using a Gaussian kernel to compute sample-wise GSVA enrichment scores for selected GO:BP terms, including cilium movement, cilium organisation, RNA splicing, and regulation of synaptic plasticity. GSVA scores were merged with sample metadata and summarised across developmental stages by age-wise means. Individual sample scores and smoothed age-dependent trends were visualised to characterise dynamic pathway behaviour across development, and gene-level coregulation profiles were generated to illustrate coordinated expression patterns within selected gene sets.

Data visualisation was performed in R mostly using ggplot2 (https://doi.org/10.1007/978-3-319-24277-4), with volcano plots generated using EnhancedVolcano (v. 1.29.1; Blighe, Rana, Lewis, 2025, doi: 10.18129/B9.bioc.EnhancedVolcano, https://bioconductor.org/packages/EnhancedVolcano; last accessed: 05 May 2026) and volcano3D^49^, and heat maps generated using ComplexHeatmap^50^.

### UMAP embeddings

To visualise the full profile of samples and their associated features, RNA-seq and protein data were Uniform Manifold Approximation and Projection (UMAP) embedded with paired plots, also visualising matching biological samples.

The top 5,000 differentially expressed genes from rlog-transformed RNA-seq data, and the full set of latent variables from the PLS-DA at the protein level, capturing major developmental trajectories, were represented as UMAP embeddings. Samples common to both datasets were connected with grey lines to illustrate the correspondence between RNA and protein measurements across conditions.

Expression of selected genes/proteins was chosen to capture the major axes of variation across developmental time and cell populations.

### Coordinated mRNA–protein dynamics analysis (<u>PLS-DA model)</u>

To identify coordinated RNA–protein variation across development, multi-omics integration was performed using supervised multiblock partial least squares discriminant analysis (PLS-DA) implemented in the mixOmics R package^51^ on 17 samples across seven developmental stages (E14–P7), comprising 4,195 matched gene–protein pairs (4,187 genes and 4,195 proteins contributing to the first two components).

PLS-DA decomposes each data omic layer into latent components defined by loading vectors (w□□□ and w□□□□), representing the contribution of individual genes and proteins to each component. Projecting each omics block onto its corresponding loading vectors yields sample-level score vectors (I□□□ and I□□□□). These scores define the latent variable structure of the integrated model and capture coordinated RNA–protein variation associated with developmental progression (LV□□□, LV□□□□) (**Fig. 3a**).

Before fitting the PLS-DA model, both omic layers were centred and variance stabilised. The weighted design parameter (λ = 0.1) controlled RNA–protein coupling strength, promoting moderate dependence between layers without enforcing complete coupling.

Developmental age was included as a categorical outcome variable to orient component extraction towards stage-discriminative variation.

The first two latent components (LV1 and LV2) were retained as they captured the principal variance of the integrated datasets and showed strong correlations between RNA and protein scores (Component 1: r = 0.957, R² = 0.916; Component 2: r = 0.866, R² = 0.75). Samples followed a developmental trajectory along these latent components, clustering into progenitor (E14–E16), migration (E17–E18), and postnatal (E19–P7) phases, with centroids and confidence ellipses highlighting coordinated trajectories across RNA and protein layers (**Fig. 3b**).

For each component, genes were ranked by the absolute magnitude of their RNA and protein loadings. Concordant genes were defined as those exceeding the 80th percentile in both layers for the same component. Concordant genes from components 1 and 2 were pooled, yielding 421 genes showing coordinated RNA–protein variation. For each concordant gene, RNA and protein expression were modelled independently as a function of developmental stage using ordinary least squares regression, with stage encoded as an ordered numeric variable from E14 to P7. The regression slope was used as an effect-size measure of temporal change. Genes were classified as concordantly upregulated (RNA and protein slopes > 0.1), concordantly downregulated (both slopes < −0.1), or non-concordant otherwise (Fig. 3d). The slope thresholds (±0.1) were selected to distinguish clear directional trends from near-flat trajectories and correspond approximately to the upper range of observed slope magnitudes (**Fig. 3d**).

For heatmap visualisation, expression values were averaged per gene and developmental stage, and z-score scaled within each gene. Genes were ordered by hierarchical clustering of scaled RNA expression profiles using Euclidean distance. Representative genes (top five per pattern, ranked by maximal absolute loading) were plotted across developmental stages using LOESS smoothing with 95% confidence intervals.

Using the same integration strategy as for the untreated condition, centred and variance stabilised RNA-seq counts and 15 matched protein abundances from pGlast+/pNeuroD+ cells were jointly analysed with multiblock PLS-DA across three developmental stages (E18, E19, P3), including both control and MIA-treated samples. Cross-omics covariance was promoted using a weighted design matrix. The analysis considered 4,217 matched gene-protein pairs and retained 4,188 RNA and 4,217 protein features in the first two components. The fitted model was used solely to create a scaffold for the protein data, which was then used to generate the UMAP embeddings in **Figure 4b**.

### Validation of RNA-seq experiments

RT-PCR was performed to validate RNA-seq results. Different aliquots of 10,000 cells from the first three experiments for timepoints E14 to E19 (E and P pooled) were reverse transcribed to cDNA with the SuperScript^TM^ IV CellsDirect^TM^ cDNA synthesis kit (ThermoFisher, #11750150). Samples of 0.5 ng/μL were used as input with a final sample volume of 20 μL. Two custom TaqMan Array Standard 96-well plates (Format 16, 6 plates each, **Supp. Fig. 1e**) were designed to enable high-resolution time-course analysis beyond standard platforms. Exon-spanning probes for twelve progenitor marker genes^52–59^ and eleven neuronal marker genes^60–70^ were selected. Samples were measured on a 7900HT Real-Time PCR System. Raw data are deposited under the GEO Accession number GSE249208.

RT-PCR data were analysed using Cq values generated by the Applied Biosystems^TM^ relative quantification analysis software module (v. 4.3). Well results, including Cq values, were transformed from raw Ct values by the latest internal algorithm of this software. Cq values above 40 were considered as not available (NA); in case of one NA, the data were replaced by the median of the other two replicates, while in case of more than one missing value, the NA was kept for analysis.

Normalisation was performed in two steps based on the ΔΔct method^71^: first, all values were normalised against the endogenous control gene of the respective panel (here: 18S (18S-Hs99999901_s1)) as a common endogenous TaqMan control. Secondly, the values were normalised to an internal plate control (here: E14 (pGlast-dsRed2/pCAG-Venus transfected)) to acquire relative quantification (RQ) values. RQ values were averaged across the three replicates of each cell population for each gene and time point by applying the geometric mean, which was the final input for the log2 transformation. RT-PCR log₂ fold changes were then plotted alongside the corresponding RNA-seq log₂ fold changes.

### DNA methylome analysis

The approach of enzymatic methyl sequencing (EM-seq)^72^ was used here to analyse 200 ng of DNA from four PolyI:C and two control samples from the pNeuroD+ cell population at E18.

The NEBNext Enzymatic Methyl-seq Kit (NEB #E7120) was used for library preparation according to the manufacturer’s protocol, applying the protocol variant optimised for large-insert libraries (option B; 359). After NEB library preparation, positive and negative spike-in controls (0.1 ng/μL of an artificially methylated pUC19 plasmid and 2 ng/μL of unmethylated λ-DNA) were diluted 1:50 and added to the samples at a volume of 1 μL per 48 μL of input DNA before fragmentation to assess the efficiency of the bisulfite conversion process. DNA libraries were analysed prior to sequencing using a High Sensitivity DNA assay on the Agilent 2100 Bioanalyzer (Agilent Technologies, #5067-4626).

The prepared libraries were sequenced on an Illumina NextSeq 2000 platform using the P3 sequencing reagents at 300 cycles in a paired-end configuration (2×151 bp plus 2×10 bp for index reads). A sequencing depth of at least 50 million read pairs per sample was targeted.

EM-seq data were first quality checked with FastQC (v. 0.11.8; Andrews et al., 2019, http://www.bioinformatics.babraham.ac.uk/projects/fastqc/; last accessed: 09 March 2026). All samples met the typical criteria for EM-seq data, i.e. a low C content, a high T content and an average (25 %) A and G content. Illumina adapters were trimmed using TrimGalore! with cutadapt (v. 3.1; 10.14806/ej.17.1.200) and aligned against the integrated methylated mm10 reference genome of bwameth (v. 0.2.6; https://doi.org/10.48550/arxiv.1401.1129; last accessed: 05 May 2026). The Picard tool MarkDuplicates (v. 2.18.2; http://broadinstitute.github.io/picard/; RRID: SCR_006525; last accessed: 05 May 2026) was then used with default parameters to remove PCR duplicates. Methylation metrics per base were extracted and converted to a MethylKit-compliant format (--methylKit) using MethylDackel (v. 0.5.2; https://github.com/dpryan79/MethylDackel.git; last accessed: 06 May 2026) with the mm10 reference genome. The resulting data matrix was then analysed using the R package methylKit (v. 1.26.0; doi: 10.1186/gb1120121113111011r87).

The Spike-in controls (pUC19 and λ-DNA) confirmed sequencing quality and the use of the pNeuroD1 construct for transfection (see **Supp. Fig. 9a**). Sequencing coverage and library complexity were comparable between treated and control samples, with similar numbers of mapped reads and median CpG coverage (**Supp. Fig. 9b**). Variation in log₁₀-transformed CpG coverage across samples showed no association with treatment or batch, indicating consistent sequencing quality across all conditions (**Supp. Fig. 9c**).

## Data availability

Raw RNAseq data are available in the NCBI Gene Expression Omnibus platform (Accession number GEO: GSE248250) together with the RT-PCR data under accession number GSE249208.

The mass spectrometry data have been deposited to the ProteomeXchange Consortium via the PRIDE^73^ partner repository with the dataset identifier PXD046067. The data were sorted according to the Sum PosteriorErrorProbability (PEP) score, which evaluated the validity of the protein detection – a high score meant high validity.

The EM-seq data is available upon request.

Data supporting the findings described in this manuscript, as well as custom code for analyses and figures, are provided in the Supplementary Information and are available from the corresponding author upon request. The code will be made publicly available upon journal publication.

### Immunofluorescence

Embryonal brains were fixed in 4% paraformaldehyde solution at 4°C overnight. The next day, the solution was exchanged with 30% sucrose in 1x PBS until the brains fell to the bottom of the tube due to water loss. Fixed brains were frozen in Tissue-Tek^®^ O.C.T.TM Compound (#R1180, Plano), cut with a Leica CM3050S cryostat, and mounted on Menzel-Gläser (Superfrost Ultra Plus, Thermo Scientific, #J3800AMNZ).

For migration analyses, the pCAG-tDimer+ cell distribution of eight control and six embryonal brains from PolyI: C-treated mothers were analysed (**Supp. Fig. 8**).

For immunofluorescent staining, the slices were blocked with 10% donkey serum (DS, Sigma, #D9663-10ML) in 1x PBS with 0.5% Triton X-100 (Roth, #3051.2) for 3h at room temperature to prevent non-specific binding. Primary anti-SatB2 (abcam, #ab92446) antibody was incubated in 5% DS in PBS with 0.25% Triton X-100 overnight at 4°C. After three PBS washes, the slices were incubated with the respective donkey anti-rabbit Alexa Fluor 647 secondary antibody (Invitrogen; #A-31573) at room temperature for 3h, followed by three washing steps with PBS along with Hoechst dye (1:1,000, Invitrogen, #33258) to stain for DNA content in nuclei. Coverslips were mounted onto slides using Fluoromount-G (SouthernBiotech, #0100-01).

Images with and without immunofluorescent staining were acquired on a Zeiss Axio Imager 2 and LSM 800 confocal microscopes. Objectives used were Plan-Apochromat 4×/0.1, Plan-Apochromat 20×/0.8 M27 (catalog 420650-9901-000), and Plan-Apochromat Korr 40×/0.95 M27 (catalog 420660-9970-000). Image acquisition employed the Airyscan detector and ZEN 3.1 (blue edition) software. 4x Images were acquired on a Nikon TE 2000-S microscope with a CoolSNAP EZ camera (Roper Scientific) using a Plan Fluor 4×/0.13 objective (WD 17.1 mm). All images were subsequently processed with Fiji v. 2.9.0/1.54d.

## Figure captions

**Supplementary Figure 1.**
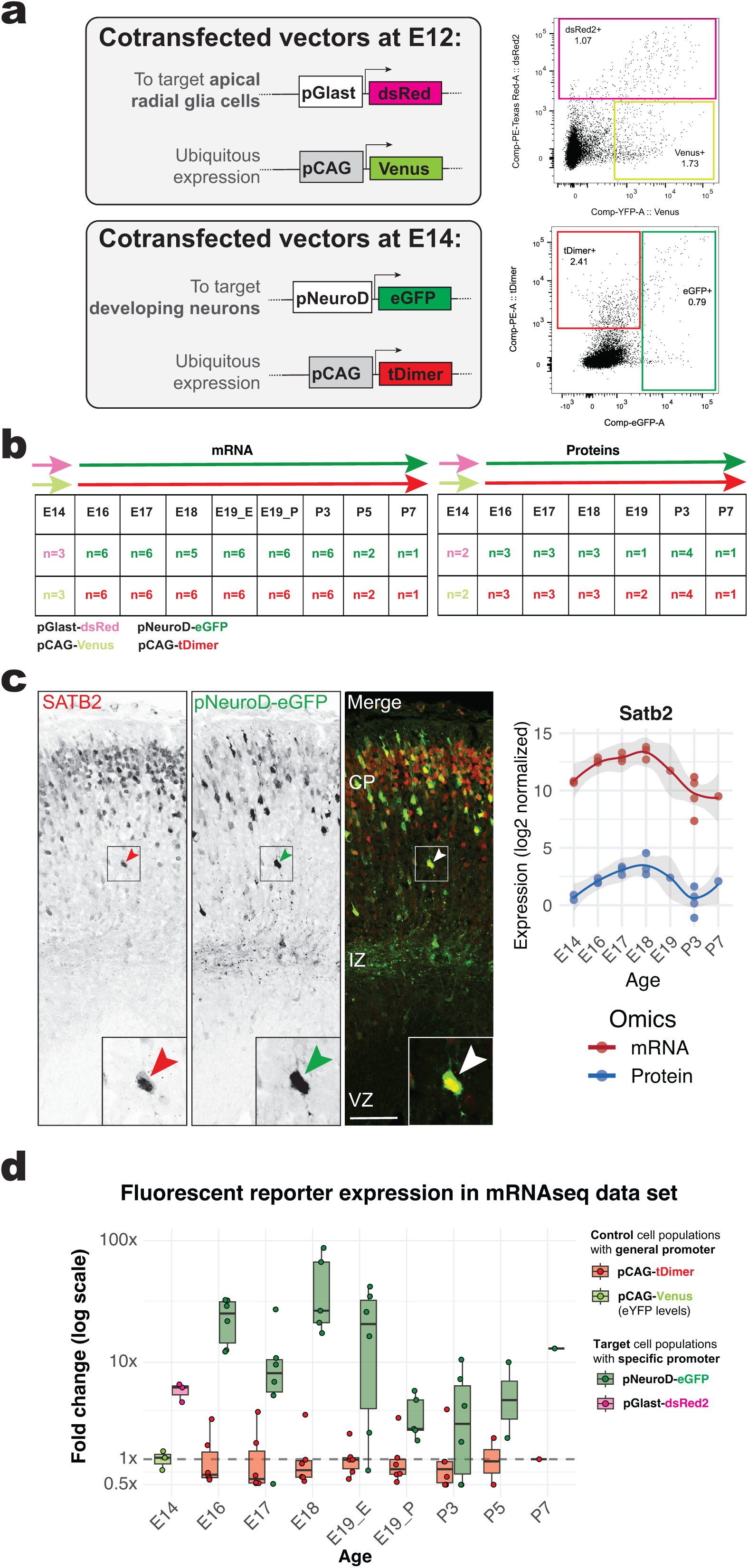
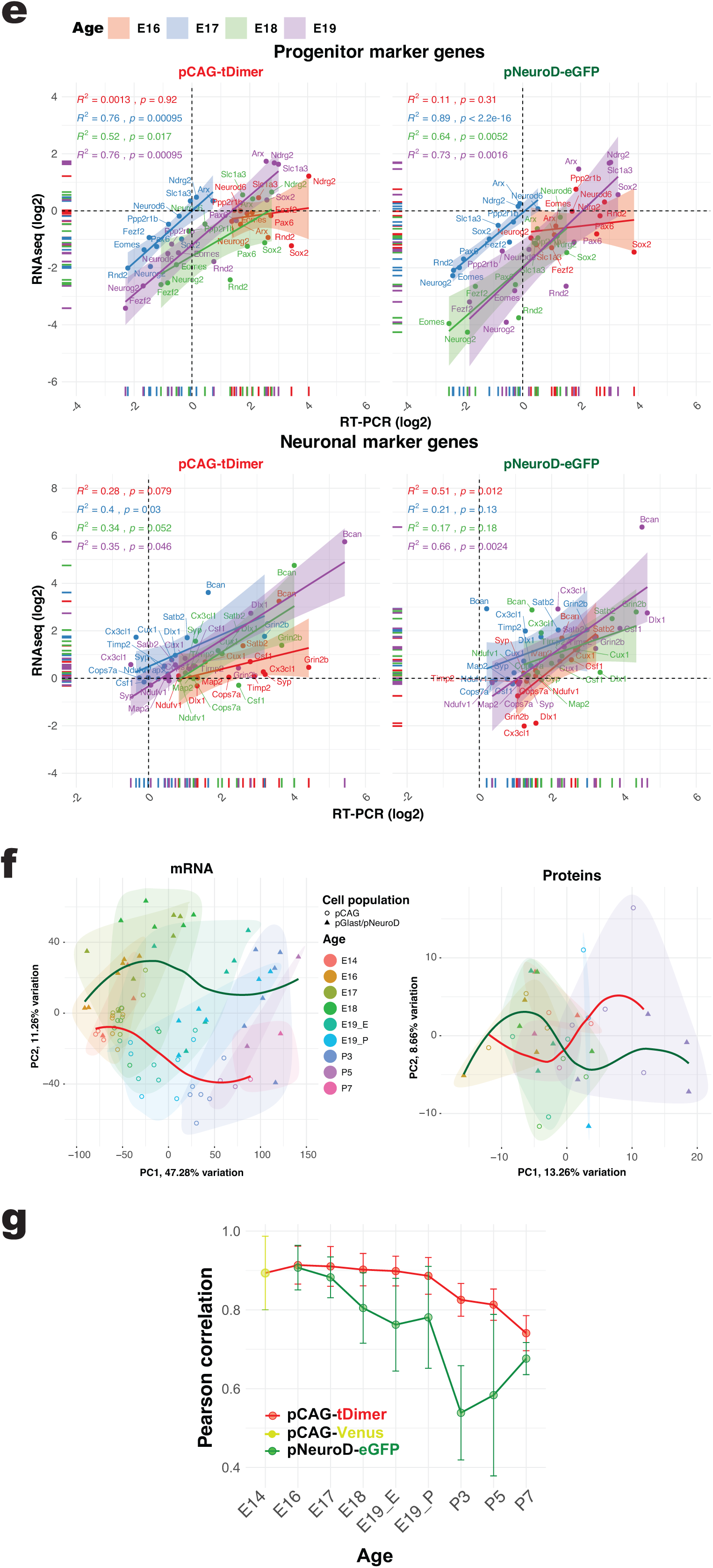
Experimental overview and validation of cell type-specific pull-down approach. **a)** *Constructs with different promoter sequences are used to target distinct cell populations.* Three promoter sequences specified cell identities and drove the expression of four fluorophores: For RGC-specific expression, pGlast was used to drive the expression of dsRed2 via IUE at E12. For expression in developing neurons, pNeuroD was used to drive expression of eGFP via IUE at E14. To target a mixed population, when the construct was introduced via IUE at E12, Venus was expressed under the CAG promoter. When the construct was introduced via IUE at E14, pCAG drove expression of tDimer. Shown adjacent are representative examples of flow cytometry gating schemes for FACS at E14 (pGlast+/Venus+) and at E18 (pNeuroD+/tDimer+). **b)** *Number of samples per developmental stage (E14–P7) and cell population.* A similar number of replicates were used for transcriptomic and proteomic profiling, and all proteomics samples had a corresponding mRNA sample. **c)** *Presence of upper-layer marker SatB2 in the analysed neuronal cell population.* Immunofluorescence staining for Satb2 shows that the overlaid pNeuroD1-eGFP+ cells were positive for Satb2 in representative coronal sections of the E18 somatosensory cortex after IUE at E14. Scale bar: 100 µm. **Right panel**: Temporal dynamics between Satb2 mRNA and corresponding protein log2 levels. **d)** *Construct mRNA abundance in pCAG and pGlast+/pNeuroD+ cell populations across developmental stages.* Boxplots display dsRed2/eGFP fold change relative to the mean pCAG expression within each experiment, with individual samples overlaid as jittered points. The dashed line indicates parity with the control expression (1×). eGFP expression in pNeuroD cells is substantially higher than in controls, often reaching one to two orders of magnitude greater, particularly at later developmental stages (see method section). The y-axis is shown on a pseudo-logarithmic scale to accommodate the wide dynamic range of expression. **e)** *RT-PCR correlates with mRNA transcript abundance detected by RNA-seq.* Spearman test confirms high correlation between RT-PCR (x-axis) and RNA-seq (y-axis) mean counts for the selected genes. Consider that the RT-PCR values shown are calculated differently from the RNA-seq values (see method section). The lines for each time point show the normalised log2 fold change of the RNA-seq data calculated by DESeq2 with contrast to E14 plotted against the log2-transformed mean of the RQ values from RT-PCR. **f)** *Principal component analysis of the untreated datasets reproduced the gradual progression of indirect neurogenesis*. PC1 showed a fan-shaped clustering of consecutive harvesting time points. PC2 explained the difference in population (“ctrl”/red LOESS curve vs. “special”/green LOESS curve). The lower 99% of variables were removed for PCA. Note that the E14 samples overlapped the pCAG-tDimer+ samples of the other prenatal time points, and all pNeuroD1+ samples clustered outwards. In contrast, the protein data did not cluster well, either by developmental stage or by cell population. The first two PCs together could only explain 21.92% of the variance in the protein data set. **g)** *Pearson correlation of E14 pGlast+ samples with all other developmental samples across ages and cell populations*. The high correlation of E16 to E18 pNeuroD+ samples with the E14 pGlast+ samples indicates that, until migration, early-born neurons retain progenitor molecular characteristics. At the postnatal stage, pNeuroD+ samples have significantly diverged from the pCAG+ cell population. Error bars show mean ± SD for cell population contrasts.

**Supplementary Figure 2.**
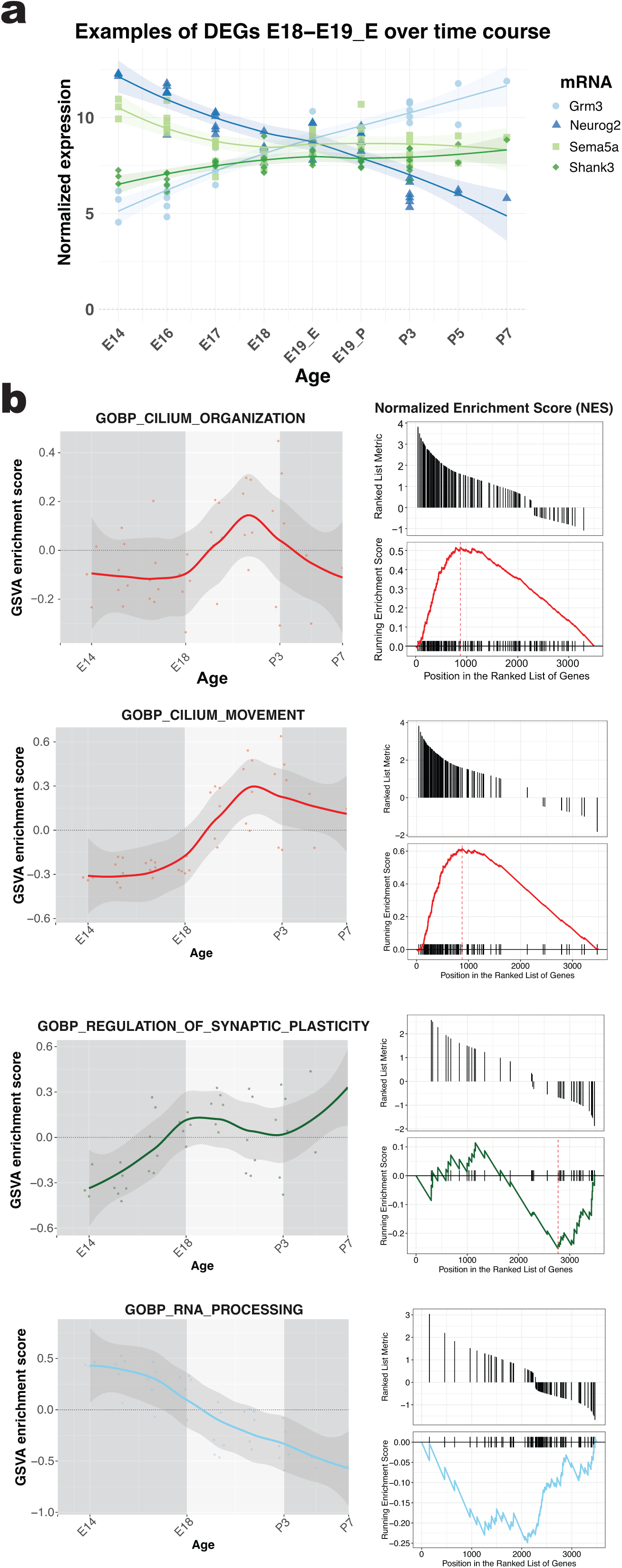
Birth associated gene set enrichment. **a**) *Representative synaptic genes show coordinated transcriptional reprogramming at birth.* Variance-stabilised RNA-seq expression of *Grm3, Neurog2, Sema5a,* and *Shank3* across development reveals opposing temporal trajectories that intersect at birth, marking a transition from prenatal to postnatal transcriptional states. Prenatally enriched genes decline after birth, while others increase postnatally, consistent with a developmental regulatory switch at birth. **b**) *Gene set enrichment and variation analyses revealed strong developmental regulation of cilium-related pathways.* **Left panels**: GSEA of cilium organisation (GO:0044782, NES = 2.03, padj = 3.35 × 10⁻¹¹) and cilium movement (GO:0003341; NES = 2.38, padj = 2.10 × 10⁻¹) showing robust positive enrichment driven by coordinated upregulation of motile cilia genes. For comparison, the most opposite trajectories among the top 5,000 DEGs (padj < 0.01) from E14 to P7 were highlighted. The most negatively enriched pathway was associated with RNA splicing via transesterification reactions (GO:0000375; NES = – 2.62, padj = 7.60 × 10⁻, on E14-P7: NES –3.47 (padj: 5.3×10-3)), indicating reduced expression of spliceosomal components. Regulation of synaptic plasticity was the most positively enriched pathway from E14-P7, even if modestly negatively enriched during the period of E18-P3 (GO:0048167; NES = –1.56, padj = 4.43 × 10⁻³, on E14-P7: NES 2.62 (padj: 3.9×10-3)). **Right panels:** GSVA profiles reveal pronounced developmental modulation of cilium-related and RNA processing pathways across embryonic and postnatal stages. GSVA applied to rlog-transformed RNA-seq data shows sample-wise pathway activity, with points indicating individual samples and smoothed curves showing age-averaged trends. Positive scores indicate coordinated pathway upregulation and negative scores indicate downregulation, highlighting pathway-specific temporal regulation during neurodevelopment.

**Supplementary Figure 3.**
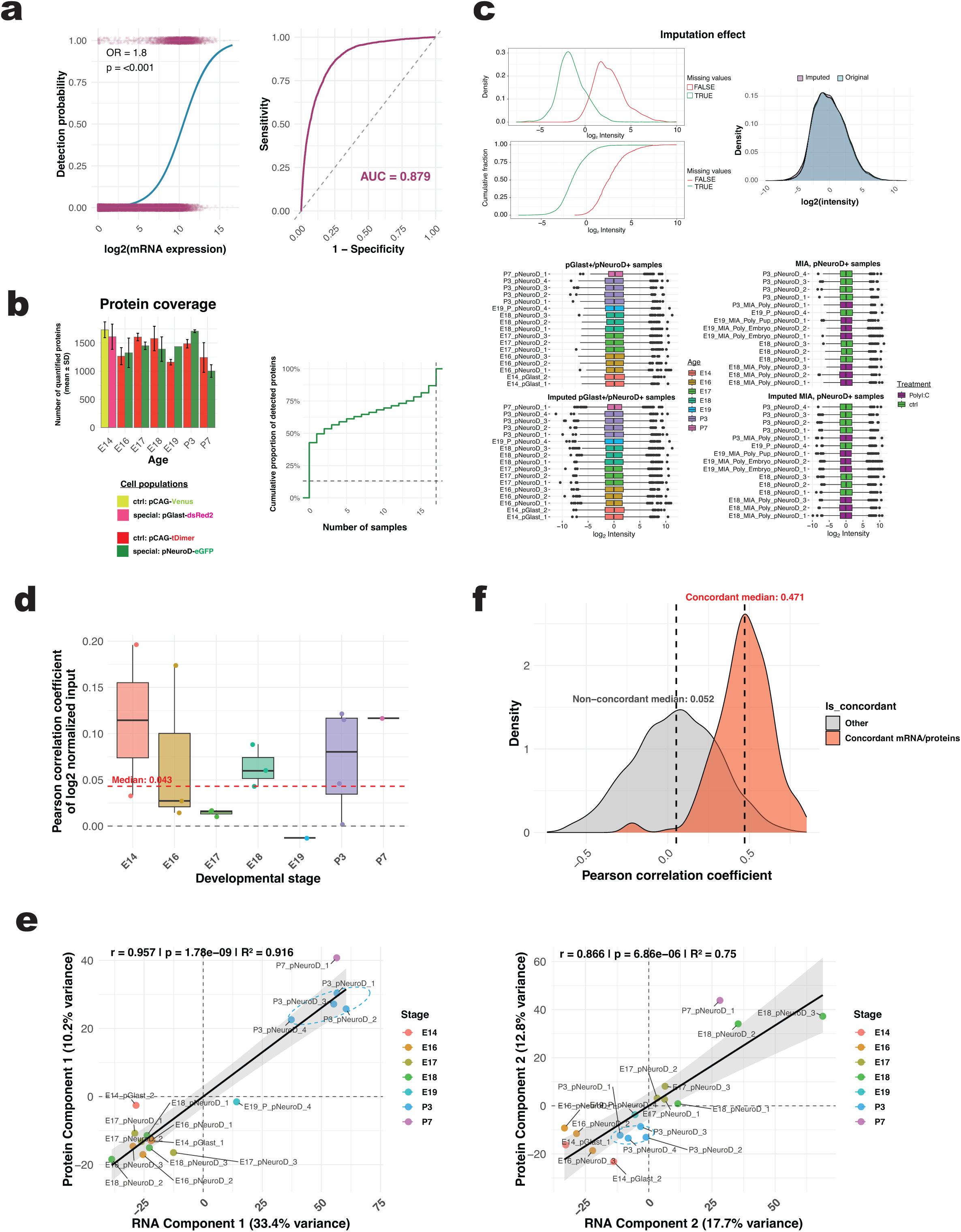
Metrics used for analysis of the proteomic data set and for integration with the transcriptomic data set. **a)** *Transcript abundance predicts protein detectability in mass spectrometry*: Each doubling of RNA expression increases the odds (OR= odds ratio) of MS detection by ∼1.8-fold (95% CI: 1.77-1.84, p < 0.001). The ribbon indicates the 95% confidence interval; points represent observed detections. ROC (=receiver operating characteristic) curve shows that RNA abundance alone predicts MS detectability well (AUC = 0.879). **b)** *The total number of quantified proteins was similar for all samples.* Regardless of age or cell population, 13.1% of proteins were consistently detected in every sample when focusing on the 17 untreated samples from E14 to P7 (similar for all conditions). **c)** *Imputation effect on detected proteins*. Proteins with missing values have, on average, low intensities (MNARs). To account for those MNARs, a manual imputation for further statistical tests was required, which was performed on a low base since the dataset was already highly normalised. **d)** *Sample-wise Pearson correlation of mRNA and protein abundance across all matched features*. With an overall median of r = 0.043, E14 and P3 samples had the most similar transcriptome and proteome levels, so genes with high/low mRNA expression also tend to have high/low protein abundance. E17 samples were closer to 0 (r = 0.0158), indicating little linear relationship between mRNA and protein levels. In contrast, the mRNA level of the one E19 sample, with a correlation of r = – 0.0127, was associated with lower protein abundance. **e)** *Integration of RNA and protein data using multiblock PLS-DA.* Scatterplots display sample scores for Component 1 (left) and Component 2 (right), coloured by developmental stage. A strong RNA-protein correlation is observed (Component 1: r = 0.957, R² = 0.916; Component 2: r = 0.866, R² = 0.75), with the variance explained by each component indicated. Linear regression (black line) illustrates the overall RNA-protein correlation. Dashed ellipses represent 75% confidence intervals per group when sufficient data points are available. **f)** *PLS-DA model identifies genes with high mRNA-protein coordination.* Concordant genes/proteins (red) display 9.1x higher overall mRNA-protein correlations than the other genes/proteins (grey), with medians indicated by dashed lines. **g)** *Developmental trajectories of representative concordant genes*. RNA (red) and protein (blue) expression trajectories across developmental stages for the top five genes per pattern (upregulated, downregulated, non-concordant), ranked by maximal absolute loading. Points represent individual samples, and solid lines indicate LOESS-smoothed trends with shaded 95% confidence intervals. Expression values are shown on a log2 scale with gene-/protein-wise z-score normalisation.

**Supplementary Figure 4.**
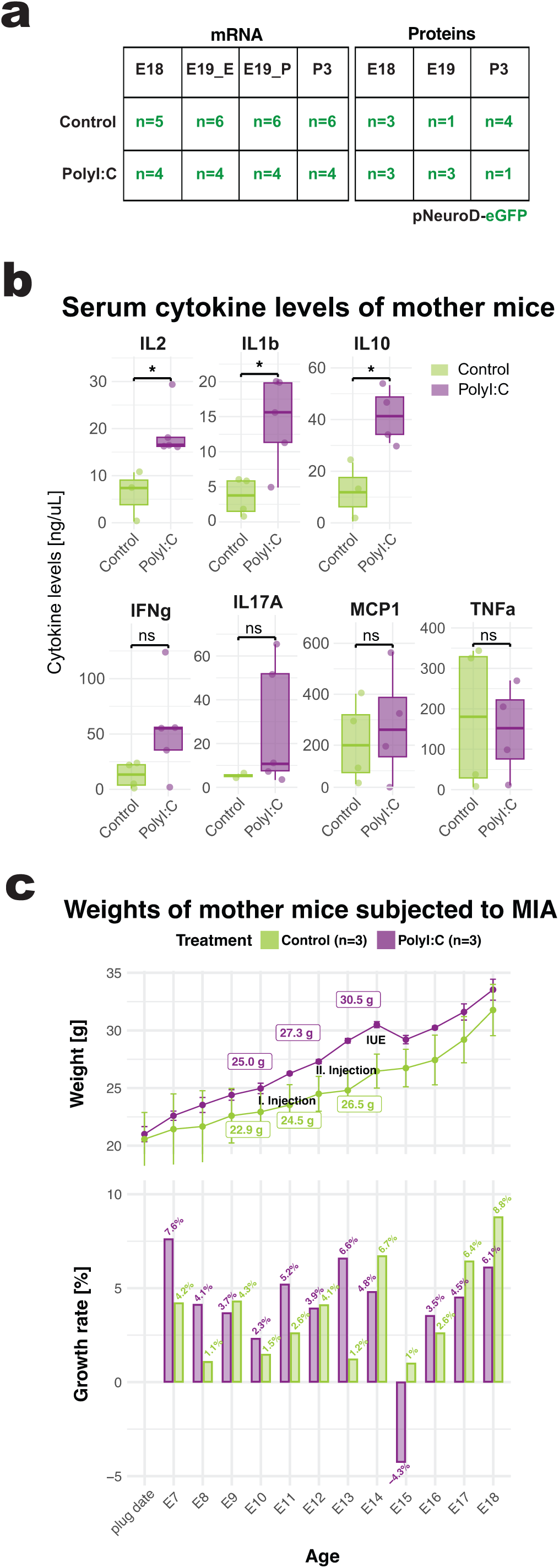
Experimental overview of the maternal immune activation approach. **a)** *Sample sizes for control and treated cohorts used for downstream analyses.* For the MIA condition, biological replicates of similar size focused on E18-P3 and the pNeuroD-eGFP+ cell population. At the mRNA level, E19 samples were distinguished as unborn embryos (E19_E) and postnatal pups (E19_P), while they were combined into a single E19 category for the proteomic analyses. **b)** *Serum cytokine levels in untreated (control) and PolyI:C-treated mother mice.* Data are presented as mean ± SEM with individual values overlaid. Statistical significance was determined by t-test for normally distributed cytokines (IFNg, IL1b, IL10, MCP1, TNFa) or Mann-Whitney test for non-normally distributed cytokines (IL2, IL17A). Points represent individual animals; bars indicate mean ± SEM; *p < 0.05, ns = not significant. **c)** *Maternal weight gain and growth dynamics following PolyI:C-induced maternal immune activation.* Mean body weights (± SD) are plotted across gestation, with growth rates (% change) overlaid as bars. Mean weights at embryonal stages (E10, E12, E14) are annotated where mice were subjected to invasive procedures such as injections and surgery. PolyI:C-treated dams were pre-selected based on their higher weight from the outset. Shown is one randomly selected experimental group with three mice per treatment; more data are available upon request. Although PolyI:C exposure was associated with a transient weight decrease after IUE at E14, the mice later recovered their growth. PolyI:C does not impair overall gestational weight gain but instead alters its temporal regulation.

**Supplementary Figure 5.**
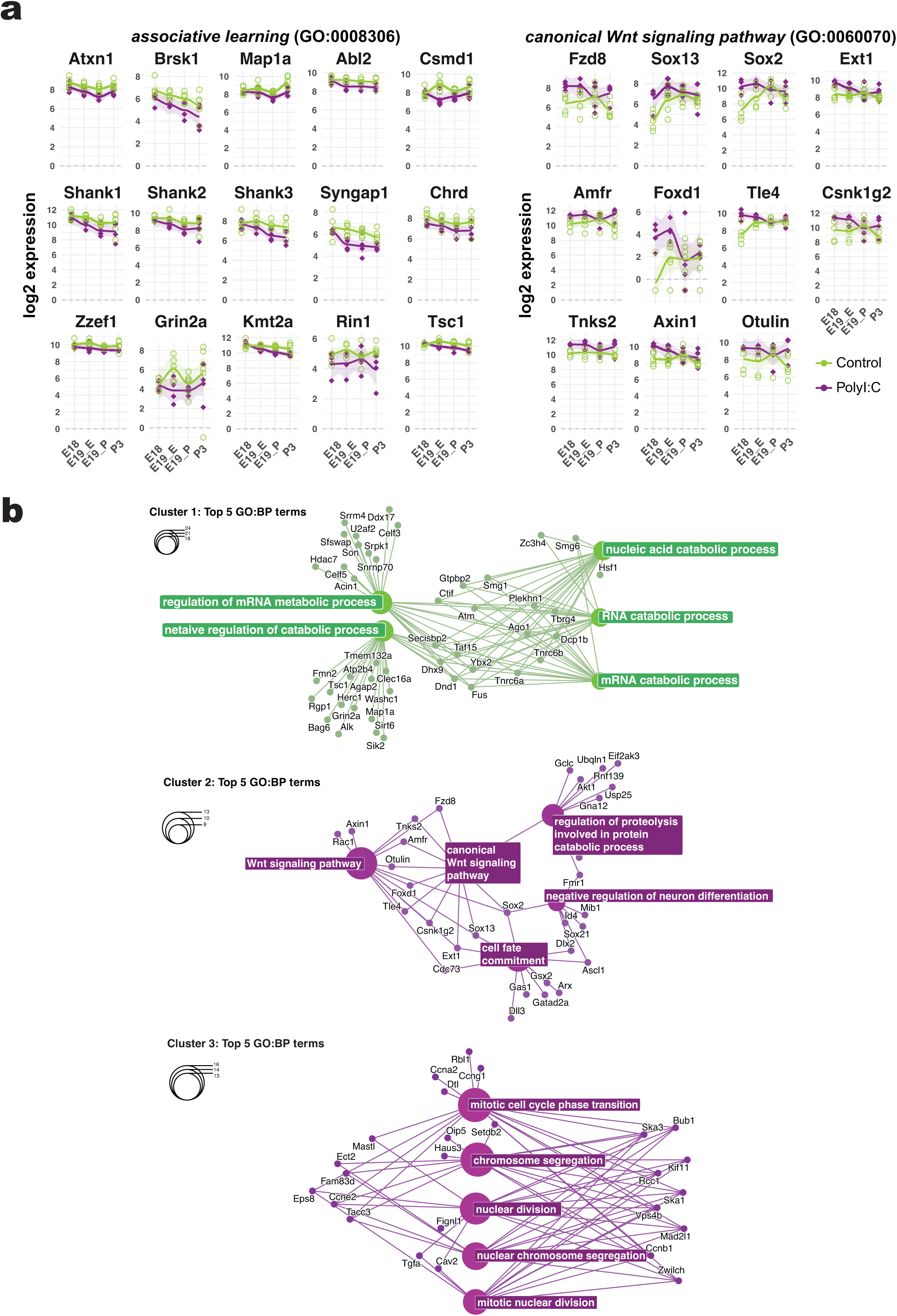
MIA treatment affects the expression of specific sets of genes. **a)** *Expression profiles of selected single genes of specific GO terms in the pNeuroD+ cell population from E18-P3 with or without MIA treatment.* Points represent individual samples, and lines indicate LOESS-smoothed trajectories for control (green) and PolyI:C-treated (magenta) animals. Expression is shown as log2-transformed counts or protein abundance, highlighting dynamic regulation across developmental time. **b)** *Overrepresentation analysis of the three clusters as cnetplots.* Clustered network plots (cnetplots) display the top five enriched GO terms for PolyI:C– and control-upregulated genes. Category nodes (GO terms) are coloured and labelled, while genes are shown as points, with genes from the corresponding gene set highlighted in black.

**Supplementary Figure 6.**
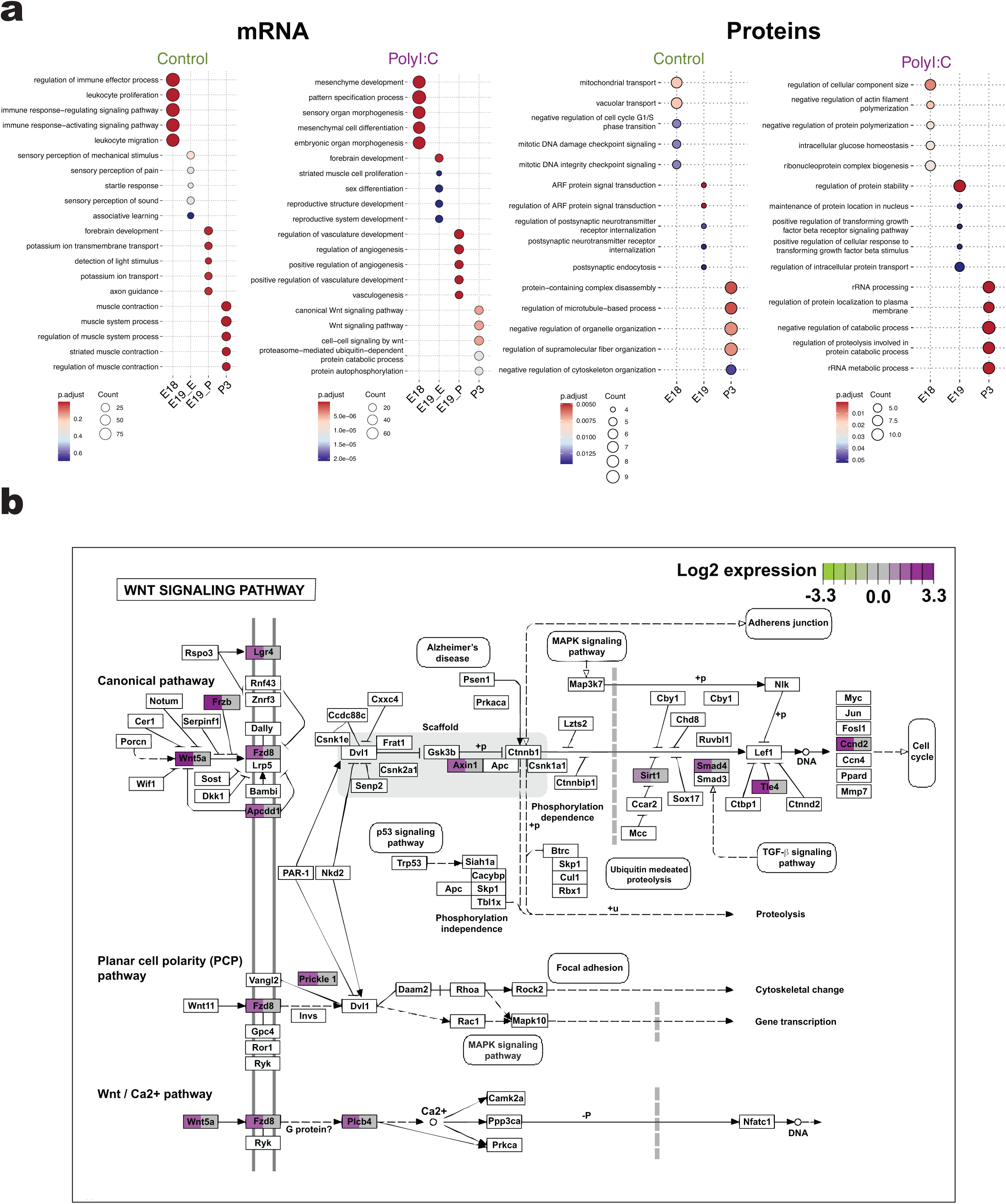
Biological processes are overrepresented in the MIA condition. **a**) Top 5 enriched GO:BP terms across consecutive time points in Ctrl (left) and PolyI:C (right) cohorts. Left panel: mRNA; right panel: proteins. At the transcript level, Ctrl samples show a developmental progression from immune activation (E18) to sensory, synaptic, and craniofacial maturation (E19E), visual/forebrain development and ion transport (E19P), and postnatal electrophysiological regulation (P3). PolyI:C samples instead enrich embryonic patterning and organ morphogenesis (E18), neural differentiation (E19_E), cell migration (E19_P), and postnatal Wnt signalling and protein regulation (P3). At the protein level, ctrl samples progress from mitochondrial and vesicle regulation (E18) to postsynaptic trafficking and RNA processing (E19), and postnatal cytoskeletal and junctional organisation (P3), whereas PolyI:C samples shift from cytoskeletal organisation and ribosome biogenesis (E18), to protein stability, trafficking, and stress signalling (E19), and postnatal RNA processing, autophagy, and protein turnover (P3). **b**) *KEGG representation of the Wnt signalling pathway (mmu04310) highlights the thirteen genes enriched in the PolyI:C cohort at E18*. PolyI:C-upregulated (log₂FC > 1) genes were identified using DESeq2 (adjusted p < 0.05) and visualised on the canonical Wnt pathway using the Pathview R package.

**Supplementary Figure 7.**
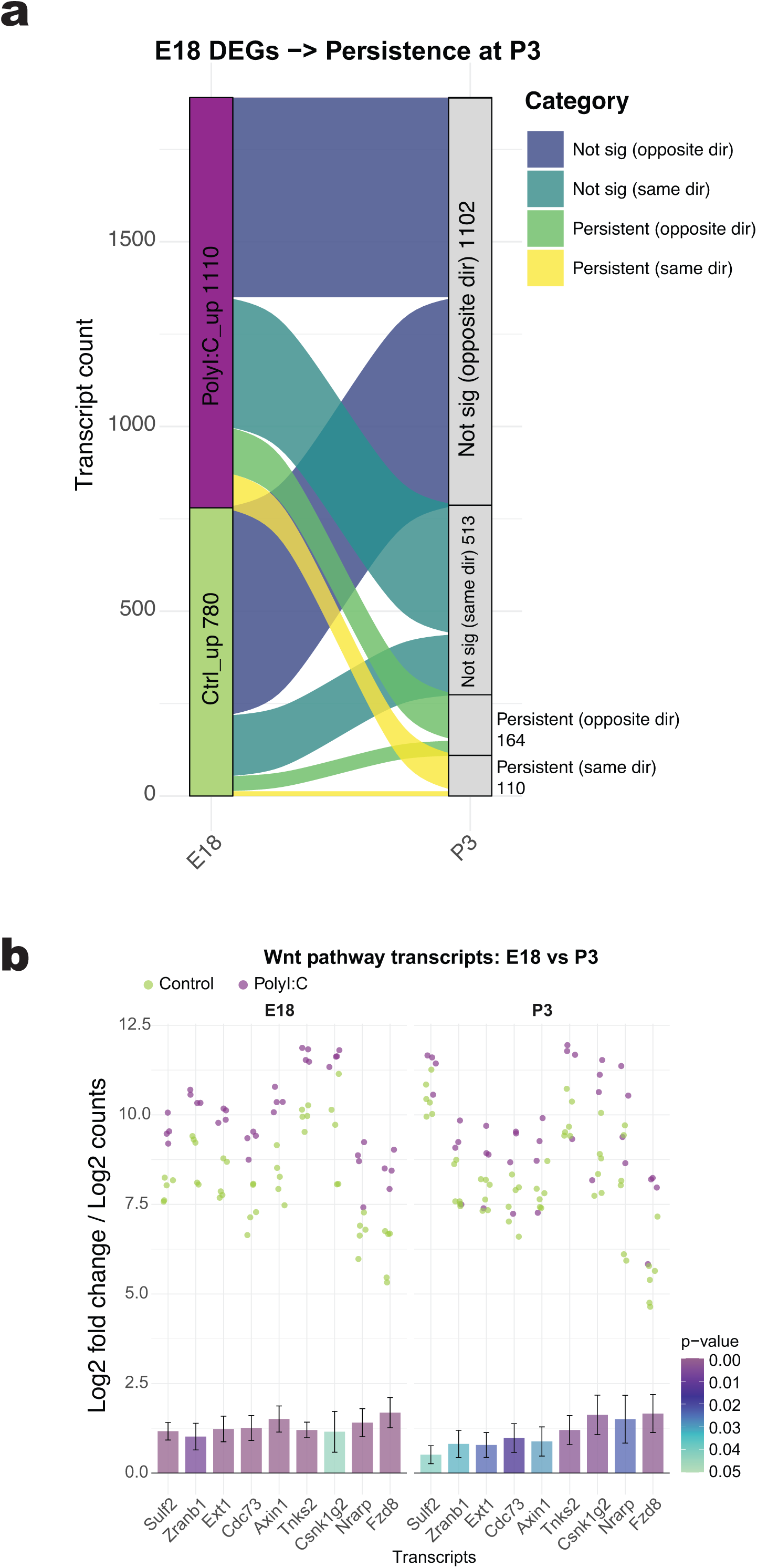
Trajectory of different gene expression after MIA treatment. **a)** *Alluvial plot tracks upregulated DEGs at embryonic day 18 (E18) to postnatal day 3 (P3).* Most PolyI:C DEGs are no longer differentially expressed at P3, but the remaining 110 genes are mainly associated with Wnt signalling, as shown in b). These genes remain upregulated at P3, although at a lower level. For example, *Fzd8* did not show a significant decrease from E18 to P3 in expression log2 level (see Supp. Fig. 5a). Sig = significant. Dir = direction. **b)** *Differential expression of Wnt pathway genes at E18 and P3.* Bar plots display DESeq2-derived log2 fold changes for PolyI:C-treated samples compared with controls at each time point. Error bars indicate the standard error of the log2 fold change (lfcSE). Bar fill denotes the nominal p-value from DESeq2 (p ≤ 0.05). Overlaid jittered points display log2-transformed normalised gene counts for individual samples, coloured by treatment group. Genes are shown side by side for E18 and P3 to highlight the persistence of directionality across developmental stages.

**Supplementary Figure 8.**
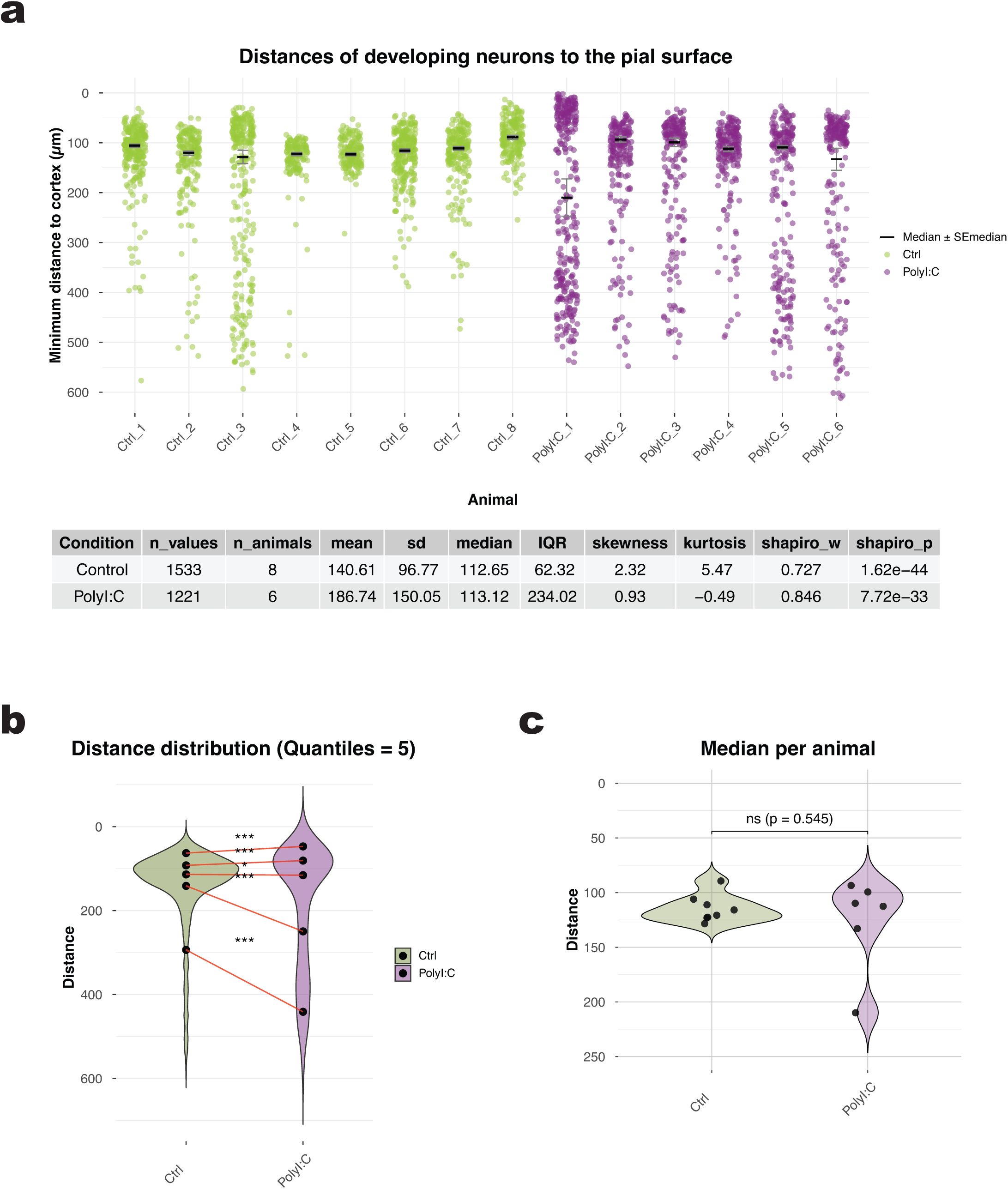
MIA treatment affects neuron position during neuronal migration in the developing cortex. **a)** *Overview of analysed cells in PolyI:C and control animals. Distance values were analysed per animal*. For each animal, the median distance was calculated, and the standard error (SE) of the median was estimated using non-parametric bootstrapping. Error bars indicate the bootstrapped SE of the median. Below: Descriptive statistics to summarise data distribution and assess normality between the control and PolyI:C-treated group. The table summarises sample size (number of values and animals), mean ± SD, median, interquartile range (IQR), skewness, kurtosis, and Shapiro-Wilk test statistics (W and p) to characterise data distribution and assess normality. Not all reported descriptive measures are shown in the graphical representation. SD, IQR, skewness, kurtosis, and the Shapiro-Wilk test consistently indicate increased dispersion and deviation from normality in the PolyI:C condition compared with the control condition. **b)** *The distribution of PolyI:C cells across cortical zones differs from that of the control.* Data were divided into quintiles based on distance (5 quantiles, 20% bins). Within each quintile, values were compared between the control and PolyI:C condition using a Poisson generalised linear model on log-transformed data with the design Distance∼Treatment. The statistical significance of the Treatment effect was assessed via an LRT against a null model without the Treatment term. P-values were corrected for multiple testing across quintiles using the FDR/Benjamini-Hochberg method (*p < 0.05; ***p < 0.001). Violin plots display the distributions with marked quantiles, revealing shifts between the control and PolyI:C condition. The slope of lines connecting quantiles indicates the magnitude of differences, with lower quantiles showing marked differences for PolyI:C cells in deeper cortical zones compared to the control. **c)** *The medians between PolyI:C and control animals are not statistically different.* Violin plots of per-animal median distances show no significant differences between conditions (standard t-test). Only the medians are compared here; all other statistical analyses in the manuscript consider the full distributions across the cortical zones.

**Supplementary Figure 9.**
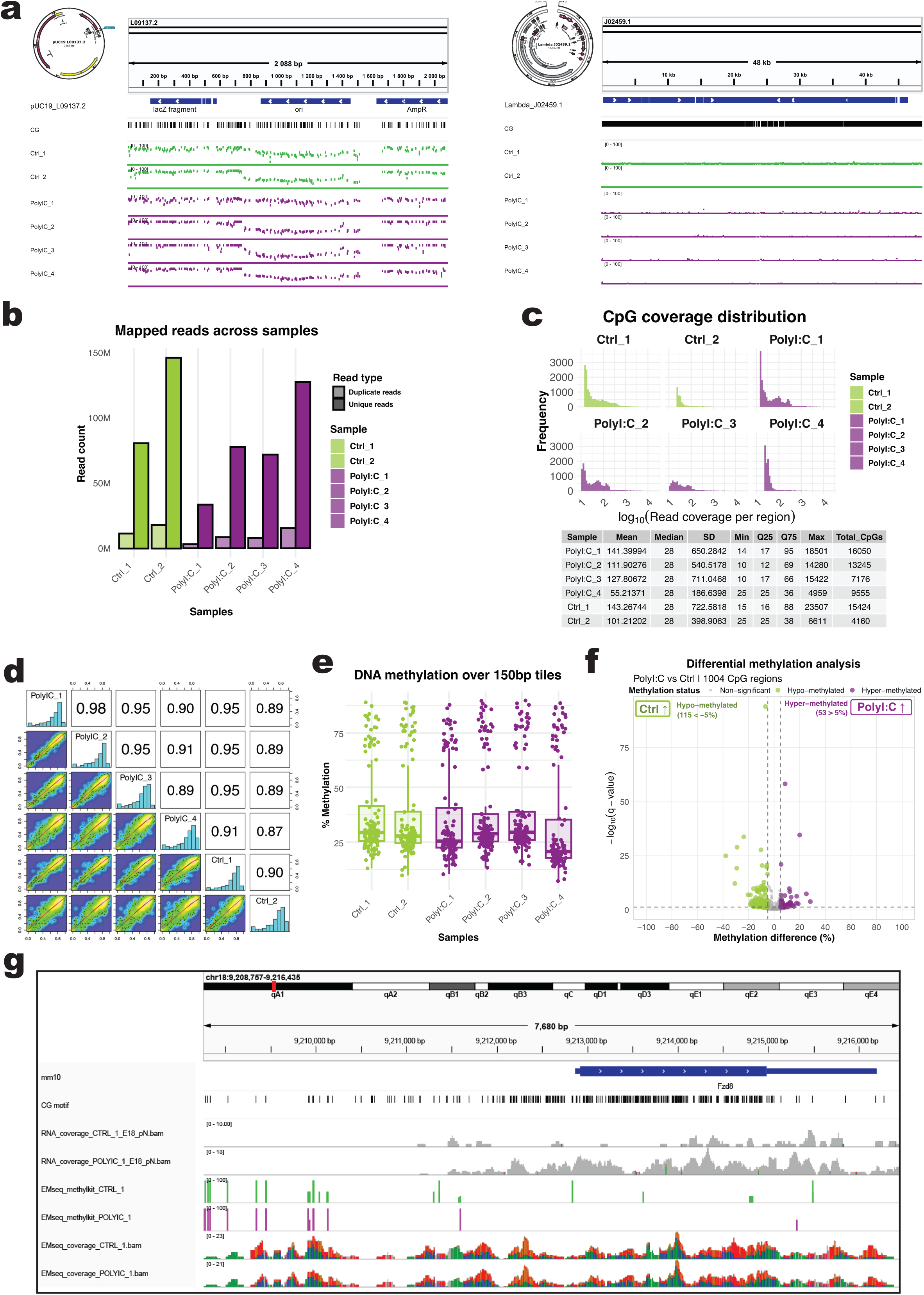
MIA treatment does not affect the methylome of upper-layer developing neurons at E18. **a)** *SpikeIn controls confirm the reliability of subsequent methylation analyses*. IGV visualisation of the lanes annotated with their respective SpikeIn controls shows that unmethylated λ-DNA (J02459.1) used as the negative control exhibited almost no methylation signal. In contrast, the artificially methylated pUC19 plasmid (L09137.2) displayed regions of 0% and 100% methylation, as well as regions with intermediate (∼50-75%) methylation corresponding to the ori and AmpR sequences of pUC19 and the pNeuroD1-eGFP construct, reflecting the co-extraction of vector-derived sequences with neuronal DNA. **b)** *Sample libraries were comparably complex.* Sample libraries were comparably complex. Bars indicate total mapped reads, partitioned into unique and duplicate reads for each sample, which behave proportionally to the overall sequenced reads, whilst the sequencing coverage/Read depths differ between samples, but are independent of PolyI:C-treated and control conditions. **c)** *CpG coverage distributions vary between samples but are not associated with condition or batch*. Below are the summary statistics of CpG read coverage (log₁₀-transformed) per sample, including mean, median, variability (SD), quartiles, and total number of covered CpG sites. 28 sequencing reads covered a CpG (median) across all samples, while total CpG counts varied slightly between libraries, as shown by right-skewed distribution tails across libraries. **d)** *Global DNA methylation is stable across samples.* Sample-wise correlation shows high pairwise correlations (Pearson r ≈ 0.87-0.98), with no condition-specific clustering. **e)** *Genome-wide DNA methylation shows no variability in DNA methylation between samples:* Boxplots show the distribution of CpG methylation levels (% methylation) across 150 bp tiles for each sample, with individual tiles overlaid as jittered points. PolyI:C-treated samples (magenta) and controls (green) show overall comparable methylation patterns. **f)** *DNA methylation differences on* the *CpG level between* the *PolyI:C and control conditions*. The volcano plot displays methylation differences (Δ%) against statistical significance (-log₁₀ q-value), with significant CpGs (|Δ| > 5%, q < 0.05) highlighted: hypomethylated (green) and hypermethylated (magenta). Dashed lines indicate significance thresholds. **g)** *Representative example of RNA-seq and EM-seq data at the candidate Fzd8 locus.* The genomic region chr18:9,209,000-9,216,500 is shown with mm10 *Fzd8* gene annotation (blue) and CpG motifs (black ticks). RNA-seq coverage tracks are shown in grey for one control and one PolyI:C sample, with lower *Fzd8* read coverage in control samples than in PolyI:C samples. EM-seq methylation tracks from methylKit are shown for a control sample (green) and a PolyI:C sample (magenta), along with coverage-normalised EM-seq reads (control, bottom coloured histogram; PolyI:C, bottom coloured histogram). Track heights indicate similar read coverage on CpG islands (CG motif in black ticks on top) for both conditions.

## References

1 Herrup, K. Post-mitotic role of the cell cycle machinery. Current opinion in cell biology 25, 711–716, doi:10.1016/j.ceb.2013.08.001 (2013).

2 Herrup, K., Neve, R., Ackerman, S. L. & Copani, A. Divide and die: cell cycle events as triggers of nerve cell death. The Journal of neuroscience: the official journal of the Society for Neuroscience 24, 9232–9239, doi:10.1523/JNEUROSCI.3347-04.2004 (2004).

3 Herrup, K. & Yang, Y. Cell cycle regulation in the postmitotic neuron: oxymoron or new biology? Nat Rev Neurosci 8, 368–378, doi:nrn2124 [pii] 10.1038/nrn2124 (2007).

4 Deneris, E. S. & Hobert, O. Maintenance of postmitotic neuronal cell identity. Nature neuroscience 17, 899–907, doi:10.1038/nn.3731 (2014).

5 Douglas, R. J. & Martin, K. A. Neuronal circuits of the neocortex. Annu Rev Neurosci 27, 419–451, doi:10.1146/annurev.neuro.27.070203.144152 (2004).

6 Velmeshev, D. et al. Single-cell genomics identifies cell type-specific molecular changes in autism. Science 364, 685–689, doi:10.1126/science.aav8130 (2019).

7 Meka, D. P. et al. Protocol for differential multi-omic analyses of distinct cell types in the mouse cerebral cortex. STAR Protoc 5, 102793, doi:10.1016/j.xpro.2023.102793 (2024).

8 Anda, F. C. et al. Cortical neurons gradually attain a post-mitotic state. Cell Res 26, 1033–1047, doi:10.1038/cr.2016.76 (2016).

9 Govindan, S., Oberst, P. & Jabaudon, D. In vivo pulse labeling of isochronic cohorts of cells in the central nervous system using FlashTag. Nat Protoc 13, 2297–2311, doi:10.1038/s41596-018-0038-1 (2018).

10 Telley, L. et al. Sequential transcriptional waves direct the differentiation of newborn neurons in the mouse neocortex. Science 351, 1443–1446, doi:10.1126/science.aad8361 (2016).

11 Stanelle-Bertram, S. et al. Male offspring born to mildly ZIKV-infected mice are at risk of developing neurocognitive disorders in adulthood. Nat Microbiol 3, 1161–1174, doi:10.1038/s41564-018-0236-1 (2018).

12 Canetta, S. E. & Brown, A. S. Prenatal Infection, Maternal Immune Activation, and Risk for Schizophrenia. Transl Neurosci 3, 320–327, doi:10.2478/s13380-012-0045-6 (2012).

13 Scola, G. & Duong, A. Prenatal maternal immune activation and brain development with relevance to psychiatric disorders. Neuroscience 346, 403–408, doi:10.1016/j.neuroscience.2017.01.033 (2017).

14 Jash, S. & Sharma, S. Pathogenic Infections during Pregnancy and the Consequences for Fetal Brain Development. Pathogens 11, doi:10.3390/pathogens11020193 (2022).

15 Hayes, L. N. et al. Prenatal immune stress blunts microglia reactivity, impairing neurocircuitry. Nature 610, 327–334, doi:10.1038/s41586-022-05274-z (2022).

16 Sabo, S. L., Lahr, J. M., Offer, M., Weekes, A. & Sceniak, M. P. GRIN2B-related neurodevelopmental disorder: current understanding of pathophysiological mechanisms. Front Synaptic Neurosci 14, 1090865, doi:10.3389/fnsyn.2022.1090865 (2022).

17 Britanova, O. et al. Satb2 is a postmitotic determinant for upper-layer neuron specification in the neocortex. Neuron 57, 378–392, doi:10.1016/j.neuron.2007.12.028 (2008).

18 Stevens, B. et al. The classical complement cascade mediates CNS synapse elimination. Cell 131, 1164–1178, doi:10.1016/j.cell.2007.10.036 (2007).

19 Zheng, S. & Black, D. L. Alternative pre-mRNA splicing in neurons: growing up and extending its reach. Trends Genet 29, 442–448, doi:10.1016/j.tig.2013.04.003 (2013).

20 Weyn-Vanhentenryck, S. M. et al. Precise temporal regulation of alternative splicing during neural development. Nat Commun 9, 2189, doi:10.1038/s41467-018-04559-0 (2018).

21 Su, C. H., D, D. & Tarn, W. Y. Alternative Splicing in Neurogenesis and Brain Development. Front Mol Biosci 5, 12, doi:10.3389/fmolb.2018.00012 (2018).

22 Furlanis, E. & Scheiffele, P. Regulation of Neuronal Differentiation, Function, and Plasticity by Alternative Splicing. Annu Rev Cell Dev Biol 34, 451–469, doi:10.1146/annurev-cellbio-100617-062826 (2018).

23 Ulicevic, J. et al. Uncovering the dynamics and consequences of RNA isoform changes during neuronal differentiation. Mol Syst Biol 20, 767–798, doi:10.1038/s44320-024-00039-4 (2024).

24 Osman, H. C. et al. Impact of maternal immune activation and sex on placental and fetal brain cytokine and gene expression profiles in a preclinical model of neurodevelopmental disorders. J Neuroinflammation 21, 118, doi:10.1186/s12974-024-03106-7 (2024).

25 Vasistha, N. A. & Sawa, A. Prenatal Immune Stress: Its Impact on Brain Development and Neuropsychiatric Disorders. Annu Rev Neurosci 48, 345–361, doi:10.1146/annurev-neuro-112723-024048 (2025).

26 Rao, Y., Wong, K., Ward, M., Jurgensen, C. & Wu, J. Y. Neuronal migration and molecular conservation with leukocyte chemotaxis. Genes Dev 16, 2973–2984, doi:10.1101/gad.1005802 (2002).

27 Lee, W. S., Lee, W. H., Bae, Y. C. & Suk, K. Axon Guidance Molecules Guiding Neuroinflammation. Exp Neurobiol 28, 311–319, doi:10.5607/en.2019.28.3.311 (2019).

28 Mikule, K., Gatlin, J. C., de la Houssaye, B. A. & Pfenninger, K. H. Growth cone collapse induced by semaphorin 3A requires 12/15-lipoxygenase. The Journal of neuroscience: the official journal of the Society for Neuroscience 22, 4932–4941, doi:10.1523/JNEUROSCI.22-12-04932.2002 (2002).

29 Kolodkin, A. L. & Ginty, D. D. Steering clear of semaphorins: neuropilins sound the retreat. Neuron 19, 1159–1162, doi:10.1016/s0896-6273(00)80408-0 (1997).

30 Klein, R. Eph/ephrin signaling in morphogenesis, neural development and plasticity. Current opinion in cell biology 16, 580–589, doi:10.1016/j.ceb.2004.07.002 (2004).

31 Sonavane, P. R. & Willert, K. Enrichment and Detection of Wnt Proteins from Cell Culture Media. Methods Mol Biol 2438, 123–131, doi:10.1007/978-1-0716-2035-9_8 (2022).

32 Knecht, S. et al. An Introduction to Analytical Challenges, Approaches, and Applications in Mass Spectrometry-Based Secretomics. Mol Cell Proteomics 22, 100636, doi:10.1016/j.mcpro.2023.100636 (2023).

33 Bocchi, R. et al. Perturbed Wnt signaling leads to neuronal migration delay, altered interhemispheric connections and impaired social behavior. Nat Commun 8, 1158, doi:10.1038/s41467-017-01046-w (2017).

34 Holt, C. E. & Schuman, E. M. The central dogma decentralized: new perspectives on RNA function and local translation in neurons. Neuron 80, 648–657, doi:10.1016/j.neuron.2013.10.036 (2013).

35 Kalish, B. T. et al. Maternal immune activation in mice disrupts proteostasis in the fetal brain. Nature neuroscience 24, 204–213, doi:10.1038/s41593-020-00762-9 (2021).

36 Choi, G. B. et al. The maternal interleukin-17a pathway in mice promotes autism-like phenotypes in offspring. Science 351, 933–939, doi:10.1126/science.aad0314 (2016).

37 Reed, M. D. et al. IL-17a promotes sociability in mouse models of neurodevelopmental disorders. Nature 577, 249–253, doi:10.1038/s41586-019-1843-6 (2020).

38 Rosshart, S. P. et al. Laboratory mice born to wild mice have natural microbiota and model human immune responses. Science 365, doi:10.1126/science.aaw4361 (2019).

39 Kanai, Y. et al. The SLC1 high-affinity glutamate and neutral amino acid transporter family. Mol Aspects Med 34, 108–120, doi:10.1016/j.mam.2013.01.001 (2013).

40 de Anda, F. C., Meletis, K., Ge, X., Rei, D. & Tsai, L. H. Centrosome motility is essential for initial axon formation in the neocortex. The Journal of neuroscience: the official journal of the Society for Neuroscience 30, 10391–10406, doi:10.1523/JNEUROSCI.0381-10.2010 (2010).

41 Niwa, H., Yamamura, K. & Miyazaki, J. Efficient selection for high-expression transfectants with a novel eukaryotic vector. Gene 108, 193–199, doi:10.1016/0378-1119(91)90434-d (1991).

42 Jacobsen, H. et al. Offspring born to influenza A virus infected pregnant mice have increased susceptibility to viral and bacterial infections in early life. Nat Commun 12, 4957, doi:10.1038/s41467-021-25220-3 (2021).

43. Voss, H., et al. HarmonizR enables data harmonization across independent proteomic datasets with appropriate handling of missing values. Nat Commun 13, 3523, doi:10.1038/s41467-022-31007-x (2022).

44 Dobin, A. et al. STAR: ultrafast universal RNA-seq aligner. Bioinformatics 29, 15–21, doi:10.1093/bioinformatics/bts635 (2013).

45 Haeussler, M. et al. The UCSC Genome Browser database: 2019 update. Nucleic Acids Res 47, D853–D858, doi:10.1093/nar/gky1095 (2019).

46 Liao, Y., Smyth, G. K. & Shi, W. featureCounts: an efficient general purpose program for assigning sequence reads to genomic features. Bioinformatics 30, 923–930, doi:10.1093/bioinformatics/btt656 (2014).

47 Love, M. I., Huber, W. & Anders, S. Moderated estimation of fold change and dispersion for RNA-seq data with DESeq2. Genome Biol 15, 550, doi:10.1186/s13059-014-0550-8 (2014).

48 Xu, S. et al. Using clusterProfiler to characterize multiomics data. Nat Protoc 19, 3292–3320, doi:10.1038/s41596-024-01020-z (2024).

49 Lewis, M. J. et al. Molecular Portraits of Early Rheumatoid Arthritis Identify Clinical and Treatment Response Phenotypes. Cell Rep 28, 2455–2470 e2455, doi:10.1016/j.celrep.2019.07.091 (2019).

50 Gu, Z., Eils, R. & Schlesner, M. Complex heatmaps reveal patterns and correlations in multidimensional genomic data. Bioinformatics 32, 2847–2849, doi:10.1093/bioinformatics/btw313 (2016).

51 Rohart, F., Gautier, B., Singh, A. & Le Cao, K. A. mixOmics: An R package for ‘omics feature selection and multiple data integration. PLoS Comput Biol 13, e1005752, doi:10.1371/journal.pcbi.1005752 (2017).

52 Arnold, S. J. et al. The T-box transcription factor Eomes/Tbr2 regulates neurogenesis in the cortical subventricular zone. Genes Dev 22, 2479–2484, doi:10.1101/gad.475408 (2008).

53 Toma, K., Kumamoto, T. & Hanashima, C. The timing of upper-layer neurogenesis is conferred by sequential derepression and negative feedback from deep-layer neurons. The Journal of neuroscience: the official journal of the Society for Neuroscience 34, 13259–13276, doi:10.1523/JNEUROSCI.2334-14.2014 (2014).

54 Bormuth, I. et al. Neuronal basic helix-loop-helix proteins Neurod2/6 regulate cortical commissure formation before midline interactions. The Journal of neuroscience: the official journal of the Society for Neuroscience 33, 641–651, doi:10.1523/JNEUROSCI.0899-12.2013 (2013).

55 Colasante, G. et al. ARX regulates cortical intermediate progenitor cell expansion and upper layer neuron formation through repression of Cdkn1c. Cereb Cortex 25, 322–335, doi:10.1093/cercor/bht222 (2015).

56 Heng, J. I. et al. Neurogenin 2 controls cortical neuron migration through regulation of Rnd2. Nature 455, 114–118, doi:10.1038/nature07198 (2008).

57 Englund, C. et al. Pax6, Tbr2, and Tbr1 are expressed sequentially by radial glia, intermediate progenitor cells, and postmitotic neurons in developing neocortex. The Journal of neuroscience: the official journal of the Society for Neuroscience 25, 247–251, doi:10.1523/JNEUROSCI.2899-04.2005 (2005).

58 Bani-Yaghoub, M. et al. Role of Sox2 in the development of the mouse neocortex. Dev Biol 295, 52–66, doi:10.1016/j.ydbio.2006.03.007 (2006).

59 Vaneynde, P., Verbinnen, I. & Janssens, V. The role of serine/threonine phosphatases in human development: Evidence from congenital disorders. Front Cell Dev Biol 10, 1030119, doi:10.3389/fcell.2022.1030119 (2022).

60 Rodriguez-Tornos, F. M. et al. Cux1 Enables Interhemispheric Connections of Layer II/III Neurons by Regulating Kv1-Dependent Firing. Neuron 89, 494–506, doi:10.1016/j.neuron.2015.12.020 (2016).

61 Nandi, S. et al. The CSF-1 receptor ligands IL-34 and CSF-1 exhibit distinct developmental brain expression patterns and regulate neural progenitor cell maintenance and maturation. Dev Biol 367, 100–113, doi:10.1016/j.ydbio.2012.03.026 (2012).

62 Schmitt, U., Tanimoto, N., Seeliger, M., Schaeffel, F. & Leube, R. E. Detection of behavioral alterations and learning deficits in mice lacking synaptophysin. Neuroscience 162, 234–243, doi:10.1016/j.neuroscience.2009.04.046 (2009).

63 Seidenbecher, C. I., Smalla, K. H., Fischer, N., Gundelfinger, E. D. & Kreutz, M. R. Brevican isoforms associate with neural membranes. J Neurochem 83, 738–746, doi:10.1046/j.1471-4159.2002.01183.x (2002).

64 Stuart, M. J., Singhal, G. & Baune, B. T. Systematic Review of the Neurobiological Relevance of Chemokines to Psychiatric Disorders. Front Cell Neurosci 9, 357, doi:10.3389/fncel.2015.00357 (2015).

65 Guo, T. et al. Dlx1/2 are Central and Essential Components in the Transcriptional Code for Generating Olfactory Bulb Interneurons. Cereb Cortex 29, 4831–4849, doi:10.1093/cercor/bhz018 (2019).

66 Bell, S. et al. Disruption of GRIN2B Impairs Differentiation in Human Neurons. Stem Cell Reports 11, 183–196, doi:10.1016/j.stemcr.2018.05.018 (2018).

67 Leone, D. P. et al. Satb2 Regulates the Differentiation of Both Callosal and Subcerebral Projection Neurons in the Developing Cerebral Cortex. Cereb Cortex 25, 3406–3419, doi:10.1093/cercor/bhu156 (2015).

68 Perez-Martinez, L. & Jaworski, D. M. Tissue inhibitor of metalloproteinase-2 promotes neuronal differentiation by acting as an anti-mitogenic signal. The Journal of neuroscience: the official journal of the Society for Neuroscience 25, 4917–4929, doi:10.1523/JNEUROSCI.5066-04.2005 (2005).

69 Marsden, K. M., Doll, T., Ferralli, J., Botteri, F. & Matus, A. Transgenic expression of embryonic MAP2 in adult mouse brain: implications for neuronal polarization. The Journal of neuroscience: the official journal of the Society for Neuroscience 16, 3265–3273, doi:10.1523/JNEUROSCI.16-10-03265.1996 (1996).

70 Chithra, Y. et al. Mitochondrial Complex I Inhibition in Dopaminergic Neurons Causes Altered Protein Profile and Protein Oxidation: Implications for Parkinson’s disease. Neurochem Res 48, 2360–2389, doi:10.1007/s11064-023-03907-x (2023).

71 Schmittgen, T. D. & Livak, K. J. Analyzing real-time PCR data by the comparative C(T) method. Nat Protoc 3, 1101–1108, doi:10.1038/nprot.2008.73 (2008).

72 Vaisvila, R. et al. Enzymatic methyl sequencing detects DNA methylation at single-base resolution from picograms of DNA. Genome Res 31, 1280–1289, doi:10.1101/gr.266551.120 (2021).

73 Perez-Riverol, Y. et al. The PRIDE database resources in 2022: a hub for mass spectrometry-based proteomics evidences. Nucleic Acids Res 50, D543–D552, doi:10.1093/nar/gkab1038 (2022).

